# PGM1 deficiency disrupts sarcomere and mitochondrial function in a stem-cell cardiomyocyte model

**DOI:** 10.1101/2025.07.01.662580

**Authors:** Silvia Radenkovic, Graeme Preston, Rohit Budhraja, Irena Muffels, Anna Ligezka, Nathan P. Staff, Ron Hrstka, Biijina Balakrishnan, Rameen Shah, Sanne Verberkmoes, Ibrahim Shammas, Inez Bosnyak, Kyle M. Stiers, Kent Lai, Lesa J. Beamer, Akhilesh Pandey, Eva Morava, Tamas Kozicz

**Affiliations:** Department of Clinical Genomics, Mayo Clinic, Rochester, MN 55905, USA; Department of Genetics, Section Metabolic Diagnostics, UMC Utrecht, Utrecht 3584 EA, NL; Department of Genetics and Genomics Sciences, Icahn School of Medicine at Mount Sinai, New York City, NY 10029, USA; Department of Laboratory Medicine and Pathology, Mayo Clinic, Rochester, MN 55905, USA; Department of Neurology, Mayo Clinic, Rochester, MN 55905; Department of Medical Genetics, University of Utah, Salt Lake City, UT 88413, USA; Manipal Academy of Higher Education (MAHE), Manipal, Karnataka 576104, India; Department of Biophysics, University of Pecs Medical School, 7624 Pecs, Hungary; Biochemistry Department, University of Missouri, Columbia, MO 65211, USA; Department of Anatomy, University of Pecs Medical School, 7624 Pecs, Hungary

**Author notes:** Authors which share the same-authorship position (Silvia Radenkovic, Graeme Preston, Eva Morava, Tamas Kozicz). Corresponding author: Tamas Kozicz, MD, PhD. And Eva Morava MD, PhD.

**Keywords:** Phosphoglucomutase-1, cardiac dysfunction, Z-disk, mitochondrial dysfunction, PGM1-CDG

## Abstract

**Background:** Phosphoglucomutase-1 (PGM1) plays a pivotal role in glycolysis, glycogen metabolism, and glycosylation. Pathogenic variants in PGM1 cause PGM1-congenital disorder of glycosylation (PGM1-CDG), a multisystem disorder with cardiac involvement. While glycosylation abnormalities in PGM1-CDG are treatable with galactose, cardiomyopathy does not improve suggesting a glycosylation-independent pathomechanism. Recently, mitochondrial abnormalities have been shown in a heart of a PGM1-deficicient patient and PGM1-mouse model. In addition, PGM1 has been associated with LDB3 (ZASP/Cypher), a sarcomeric Z-disk protein also associated with cardiomyopathy. However, the cardiac-specific role of PGM1 remains poorly understood, and targeted therapies for PGM1-related cardiomyopathy are currently lacking.

**Methods:** Induced pluripotent stem cell–derived cardiomyocytes (iCMs) were generated from PGM1-deficient patient fibroblasts. Multielectrode array (MEA) recordings, untargeted (glyco)proteomics, and pathway analysis were performed to assess functional and molecular changes. Key findings were validated using tracer metabolomics and mitochondrial respiration assays.

**Results:** PGM1-deficient iCMs exhibited reduced beating frequency, impaired contractility, and prolonged contraction kinetics. Proteomic analyses revealed depletion of Z-disk components, including LDB3. AlphaFold3 structural modeling predicted a direct interaction between PGM1 and LDB3, implicating PGM1 in Z-disk integrity, which was confirmed *in vitro*. In addition, mitochondrial proteins were severely depleted, prompting us to investigate mitochondrial function. Functional validation confirmed extensive metabolic rewiring, energy depletion, and severely impaired mitochondrial respiration. Finally, the *in silico* drug repurposing identified possible therapeutic options that could target PGM1-deficient cardiomyopathy.

**Conclusion:** PGM1 is a key regulator of cardiomyocyte function, linking sarcomeric Z-disk integrity with mitochondrial metabolism. These mechanistic insights offer a foundation for developing targeted therapies for PGM1-CDG and potentially other cardiomyopathies involving Z-disk dysfunction.

**Graphical abstract:** 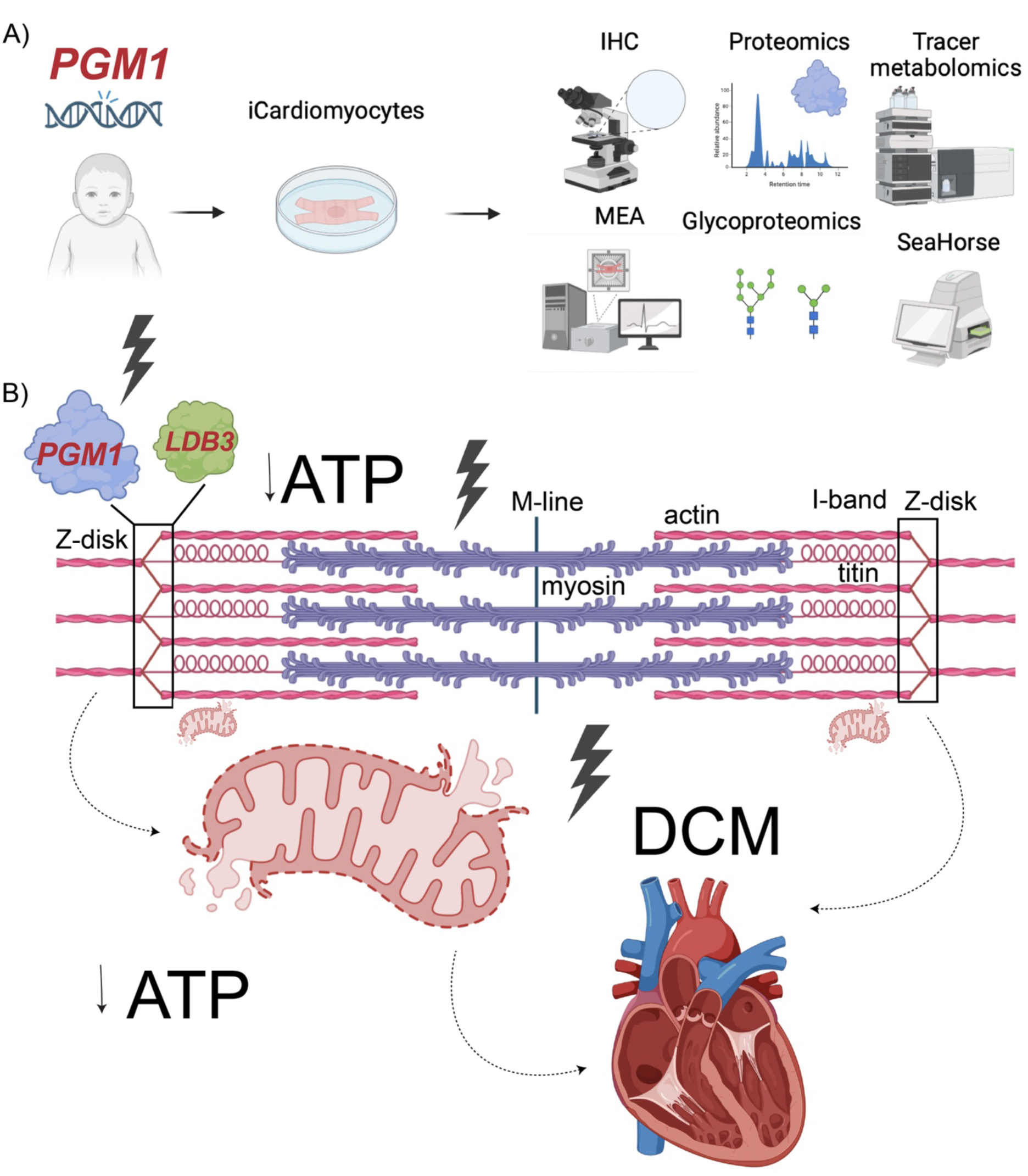

## Background

Phosphoglucomutase-1 congenital disorder of glycosylation (PGM1-CDG) (OMIM: 614921) is a multisystem disease caused by biallelic pathogenic *PGM1* variants, affecting glycogen metabolism, glycolysis, and glycosylation (**Figure (Fig) 1**) [1,2]. While clinical presentation often includes liver involvement, dysmorphic features, and coagulation abnormalities, cardiac dysfunction is the most severe, often lethal presentation in PGM1-CDG[1–4].

The age of onset of cardiac complications in PGM1-CDG is variable and can occur within the first 5 years[1,3–9]. The most frequent cardiac complication in PGM1-CDG is dilated cardiomyopathy, but enlarged left ventricle, left-ventricular non-compaction cardiomyopathy, hypertrophy, and ECG abnormalities including tachycardia and prolonged QT have also been described[3–5,7,10–12]. Cardiac failure resulting in cardiac arrest and early death was reported in more than 12% of previously reported cases[10].

Oral galactose supplementation improves glycosylation and clinical symptoms associated with glycosylation abnormalities in PGM1-CDG such as coagulation and liver function[1,3,5,11,13,14]. Unfortunately, galactose does not ameliorate cardiac presentation, suggesting that additional pathogenic mechanisms apart from impaired glycosylation are involved in cardiac dysfunction[3,5,11,12,14].

Cardiac presentation is uncommon in other CDG[4,15], however, it has been reported in almost 50% of PGM1-CDG individuals. The genotype-phenotype correlation in PGM1 deficiency was partially elucidated by the identification of two PGM1 isoforms: PGM1-1 and PGM1-2[16][1–4]. PGM1-1 is ubiquitously expressed, while PGM1-2 is specifically expressed in muscle and heart tissue [10]. The patients whose pathogenic variants affect PGM1-2 isoform have the highest occurrence of muscle and cardiac presentation[10], suggesting PGM1-2 plays a crucial role in the heart. PGM1-2 also features a unique binding motif, in comparison to PGM1-1[10]. While, no cardiac-specific PGM1 binding partner was so far reported, PGM1 has been linked to LDB3[17], a key component of the sarcomeric Z-disk[17,18]. PGM1 was first identified as a potential biding partner of LDB3 in a yeast screen[17]. In addition, immunohistochemistry in rat cardiomyocytes showed PGM1 colocalizes with LDB3 in the sarcomere, while mutant LDB3 showed reduced binding to PGM1 resulting in sarcomere disruption [17]. Since, pathogenic *LDB3* variants lead to sarcomere disruption and dilated cardiomyopathy[17,18], the role of PGM1 in LDB3 cardiomyopathy was suggested[17]. However, the role of PGM1 in cardiac function remains poorly understood.

The role of mitochondria in PGM1 cardiomyopathy has been previously speculated as abnormal mitochondria have been found in a heart of a one-year-old PGM1-CDG patient with severe cardiomyopathy, who underwent a heart transplantation[19].To further investigate PGM1-related cardiomyopathy, a cardiac specific PGM1 knock-out (KO) mouse model was developed[19]. While the model showed similar mitochondrial abnormalities on histology, the specific mechanism linking PGM1 to cardiac and mitochondrial function was not clear[19].

To elucidate the role of PGM1 in cardiac function, we leveraged multi-omics and functional analyses in patient-derived induced pluripotent stem cell derived cardiomyocytes (iCMs). PGM1-deficient iCMs exhibited reduced beating frequency, impaired contractility, and prolonged contraction kinetics, indicative of defective excitation-contraction coupling. Proteomic analysis revealed a global reduction in sarcomeric proteins, including LDB3, with AlphaFold3 modeling suggesting a direct interaction between PGM1 and LDB3, linking PGM1 to Z-disk integrity. The interaction of PGM1 and LDB3 was further confirmed *in vitro,* confirming previous findings. Apart from depletion in sarcomeric proteins, PGM1-deficient iCMs showed depletion of MitoCarta proteins. To confirm the omics findings, functional tests were performed. Tracer metabolomics showed metabolic rewiring resulting in disruption of energy metabolism and depletion of ATP. Respiration studies using Seahorse showed decreased basal respiration and ATP-linked respiration confirming mitochondrial dysfunction. As currently there are no approved treatments for mitochondrial dysfunction, *in silico* drug repurposing was used to identify potential therapeutic strategies for PGM1-related cardiomyopathy.

These findings establish PGM1 as a critical regulator of cardiomyocyte function, potentially linking Z-disk stability to mitochondrial function. By elucidating novel mechanisms underlying cardiac pathology in PGM1-CDG, this study also provides a foundation for targeted therapeutic interventions.

## Methods

### Cell culture of patient-derived fibroblasts

Fibroblasts from four patients were collected as part of clinical care via skin-punch biopsy and residual samples were stored in the Mayo Clinic FCDGC biobank (Mayo IRB: *16-004682*). Informed consent was obtained and recorded. Fibroblasts were maintained in MEM medium (Gibco, 11095080) supplemented with 10% FBS (Cardinal Healthcare M7201-127) and 1% Anti-Anti (Gibco, 15240062) in the incubator at 37 °C, 5 % CO_2_. Routine mycoplasma testing was performed.

### Generation and maintenance of human induced pluripotent stem cells (hiPSC)

hiPSCs were generated from patient fibroblasts following standardized, previously reported methods[20]. Briefly, Sendai virus Cytotune 2.0 kit (ThermoFisher) was used to generate PGM1-CDG (P1, P2, P3, P4) and healthy control hiPSC (GM8399, GM1651 Coriell Institute for Medical Research, NJ, USA). Absence of chromosomal abnormalities was confirmed by karyotype G-banding. Markers of pluripotency were assessed by flow cytometry, immunohistochemistry (Oct 4, SSEA, Nanog, Tra-1-60), and three-germ layer (trilineage) differentiation. Mycoplasma testing was routinely performed. Only the hiPSC clones meeting all the criteria were selected. All hiPSCs were maintained in mTesr Plus medium supplemented with 10% mTeSR supplement and 1% Anti-Anti (Gibco) on 60mm dishes coated with 1 mg/ml Geltrex matrix (ThermoFisher, A1413302) in the incubator at 37 °C, 5 % CO_2_.

### Differentiation of hiPSC into iCMs

Differentiation of hiPSC into the hiPSC-derived cardiomyocytes (iCMs) was performed based on the previously described chemically-defined cardiomyocyte differentiation protocol[21,22]. The protocol was optimized for each hiPSC cell line, to ensure the optimal generation of cardiomyocytes. The outline of the protocol is shown in Figure 1. Briefly, hiPSC were maintained in 60mm dishes in 10% mTeSR supplement (Stem Cell Technologies) and 1% Anti-Anti (Gibco). One confluent, hiPSC were seeded into 6-well plates coated with 1mg/ml Geltrex (ThermoFisher, A1413302) in mTeSR plus medium supplemented with 10µM ROCK inhibitor (Y-27632, Torcis 5148). Once the cells reached 90-95% confluence, they were treated with CHIR99021 (Tocris, 4423) in RPMI 1640 (Gibco, 11875093) supplemented with B27-insulin (Thermo Fisher Scientific, A1895601) (referred to as RPMI-insulin) for three days (Day0-Day2). The concentration was optimized for each cell line and between 3-8 µM was used. Next, the medium with CHIR was removed, the cells were washed with DPBS (Gibco) and the cells were treated with 5µM IWP2 in RPMI-insulin for 48h (day 3-5). After IWP2 treatment, the medium was removed, the cells washed with DPBS and new RPMI-insulin medium added every 48h, until beating is observed (usually between D7-D14). The cells that did not start beating after two weeks in culture were discarded. Once the beating was observed, RPMI 1640 medium supplemented with B27 with insulin (ThermoFisher, 17504044) (from here referred to as RPMI+insulin) was added to the cells. To purify the cardiomyocyte culture and remove other cell types, lactate selection was performed by incubating cells with RPMI 1640 without glucose (Gibco, 11879020), supplemented with Sodium DL-Lactate Solution (Sigma, L4263), Recombinant Human Albumin (Sigma, A9731) and L-ascorbic Acid (Sigma, A8960) until majority of the cells that survived are beating (1-3 days). The medium was then switched back to RPMI+ insulin for maintenance. Finally, the cells were dissociated from the plates with TrypLE (Thermo Fisher Scientific, A1217703), collected in RPMI+insulin supplemented with 10% Knock-Out Serum (KOSR, ThermoFisher 10829018) and 10 µM ROCK inhibitor Y-27632. The cells were counted and pelleted at 800 rpm, 6 min, room temperature (RT) before freezing them in 90%, 10% DMSO (Sigma, D2650), 10 µM ROCK inhibitor. Approximately 300microL of freezing medium was used for each million of cells frozen. For all the subsequent experiments, the cells were thawed in RPMI+ insulin supplemented with 10% Knock-Out Serum (KOSR, ThermoFisher 10829018) and 10 µM ROCK inhibitor Y-27632 (1:10), counted and seeded for specific experiments (see below). The viability after thawing ranged from 70-90%.

**Figure 1.**
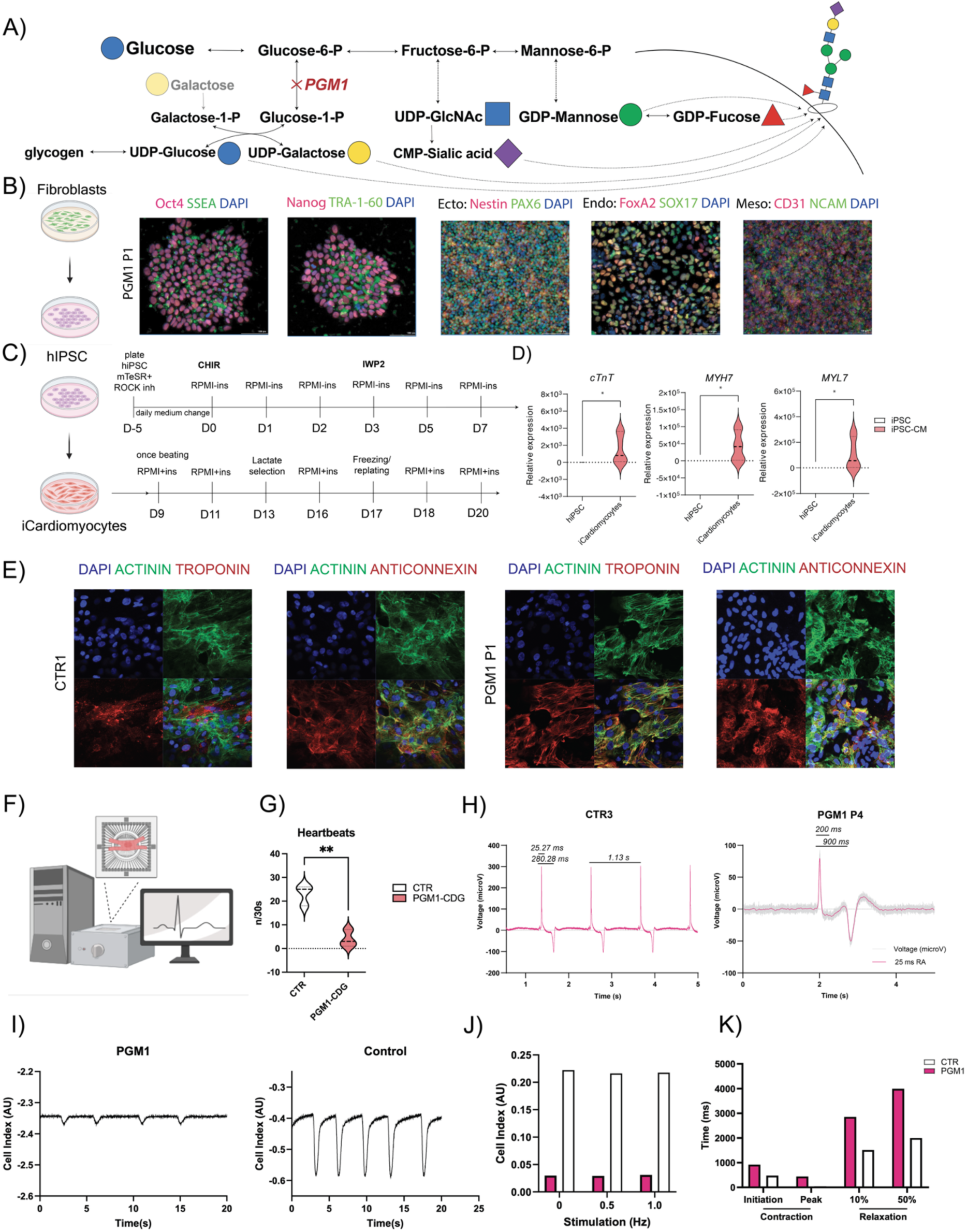
Generation and validation of iCMs (iCM). B) The successful reprogramming of fibroblasts into human induced pluripotent stem cells (hiPSC) was validated by pluripotency markers (Oct4, SSEA, Nanog, TRA-1-60) and the three-germ line differentiation: ectoderm (ecto-Nestin, PAX6), endoderm (endo-FoxA2, SOX17), and mesoderm (meso-CD31, NCAM). C) hiPSC were differentiated into iCM following the chemically defined protocol. Scale bar shows 100 μm. Cardiomyocyte markers were assessed in control and PGM1 iCM by D) RT-qPCR for cardiac troponin (*cTnT*), myosin heavy chain 7 (*MYH7)* and myosin light chain 7 (*MYL7);* and E) Immunohistochemistry (IHC) for actinin (green), cardiac troponin (red), and anti-connexin43 (anticonnexin; red). Scale bar shows 100 μm. F) Schematic representation multielectrode array (MEA) of iCM; G) PGM1-deficient iCM show decreased beating rate compared to healthy CTR. PGM1-iCM n=3, CTR iCM n=3. H) Representative MEA readouts for healthy CTR (top) and PGM1-deficient iCM. X-axis shows the time of recording in seconds (s). Y-axis shows the voltage (µV). I) PGM1 iCMs displayed a profound reduction in contractility relative to the control J) Stimulation with 0.3 ms pulses of 1200 mV at either 0.5 or 1.0 Hz in PGM1 and CTR iCM did not show any improvement in contraction amplitude. K). PGM1 iCMs displayed increased time between application of the stimulation and both contraction initiation and peak contraction. PGM1-iCMs also displayed increased time between peak contraction and both 10% and 50% relaxation. SD is shown; p value marked with * corresponds to p<0.05; ** corresponds to p<0.001.

### Immunohistochemistry

German 2-well chamber slide systems (ThermoFisher Scientific, 154852) were coated with 1mg/ml Geltrex. iCMs were thawed as described above and approximately 2 million cells were plated in each well (day 0). The following day, cells were washed with DPBS and fresh RPMI+ medium added. The medium change was performed every two days. The beating was observed in all cells after 5 days in culture. On day 7 of the experiment, iCMs were washed with 3 times with TBS (Biorad, 1706435), fixed with 4% paraformaldehyde (Sigma, 162048) for 15 minutes at RT, and rinsed with 3xTBS. The cells were then incubated for 30 min RT in blocking solution containing 0.5 % Triton X-100 (Roche, 10789704001) plus 6% goat serum (Invitrogen, 31873) in TBS. Then, the blocking solution was removed and the cell incubated overnight at 4°C in primary antibodies diluted in the blocking solution (actinin mouse, ThermoFisher A7811, 1:200, cTnT (R) (Abcam ab45932) 1:200, alpha-connexin43 (Sigma C6219) 1:600). Incubate at 4 degrees C overnight. The following day, the cells were washed 3x TBS and then incubated with the secondary antibodies (1:400, anti-mouse IgG-AlexaFluor488, ThermoFisher A28175), anti-rabbit IgG-AlexaFluor 555, ThermoFisher, A31572) at 37°C in the dark for 1h. Next, the cells were washed 3X TBS, and the TBS aspirated completely before removing the chamber sides. Gold anti-fade reagent with DAPI was added to the glass bottom and topped with a glass coverslide before sealing. The cells were visualized by confocal microscope using appropriate lasers.

### Quantitative PCR analysis to assess cardiac-specific marker expression

Briefly, 4 million iCMs were plated in RPMI+ insulin supplemented with 10% KOSR and 10µM ROCK inhibitor in 6-well plates. The following day, the cells were washed with DPBS and the medium refreshed. The medium was refreshed every 48h, for seven days. To isolate the RNA, the cells were dissociated from the plates with TrypLE (Thermo Fisher Scientific, A1217703), and collected in RPMI+insulin supplemented with 10% Knock-Out Serum (KOSR, ThermoFisher 10829018) and 10 µM ROCK inhibitor Y-27632. Then, the cells were centrifuged at 800 rpm, 6 min, RT. The supernatant was aspirated and the cells washed with 1mL DPBS, centrifuged again at 1500 rpm, 5 min, RT and the pellet snap frozen on dry ice and placed in −80°C freezer. For RNA isoltation from hiPSC, once hiPSC reached 80-90 % confluence in 60 mm plates, cells were washed with 3 ml DPBS, then 1ml of ReleSR was added. ReleSR was aspirated and the cells incubated for 5 min at 37 degrees. Next, the cells were harvested in 2mL mTeSR plus, centrifuged at 800 rpm, 6 min, RT. The supernatant was aspirated and the cell pellet washed with 1mL DPBS. The cells were centrifuged again at 1500 rpm, 5 min, RT, the supernatant was aspirated again and the cells snap frozen on dry ice and placed in −80°C freezer.

RNA mini plus isolation kit (Quiagen, 74134) was used to isolate RNA from the cell pellets and the RNA purity and concentration assessed by Nanodrop Spectrometer (ThermoFisher). Primer mix was prepared for the genes of interest (*cTNT, MYH7, MYL7, IDT*) and housekeeping gene (*ACTB, 18SRNA IDT*) by adding 1 μL of forward and reverse primer, 5 μL SYBR universal PCR master mix buffer (Applied biosciences), and 2 μL of RNase-free water to 1 μL of cDNA. The samples were transferred to a 324-well PCR plate, the plate was sealed and briefly centrifuged before running the assay on the Lightcycler RT-PCR system (Roche). Following protocol was used: 1) preincubation at 95°C for 5 min; 2) 45 amplification cycles at 95 °C, hold 10s; 60 °C, hold 10 s, 72 °C, hold 10 s; 3) melting curve 95 °C, hold 5s, 65°C, hold, 60 s, 97°C, continuous; 4) cooling at 40°C, hold 30 s. . The melt curve analysis was performed and the Ct values were exported from the program and analyzed. The 2ct method was used to analyze the relative changes in gene expression normalized against house-keeping gene mRNA expression[23]

### Multi-Electrode array (MEA)

Multi-electrode array plates (MEA, Multichannel systems) were coated with fibronectin and incubated for 1h at 37°C. Then, 350.000 cells per well were plated in RPMI+ medium. The cells were left to incubate at RT for 1h, before transferring them to the incubator, to ensure optimal seeding. After 24h, the cells were observed and the medium was changed, after which the medium was changed every 48h and iCM observed for beating daily. Once the beating was observed, MEA plates were transferred to the multi-well MEA system (Multichannel systems), kept at 37°C. The signal was recorded for 2min using MEA multichannel-screen software (V 2.20.9, Multichannel systems). The data from multiple wells was analyzed using MEA multichannel analyzer software (V 2.0.6.0 Multichannel systems).

### iCardiomyocyte Contractility

iCM contractility was assessed using the xCELLigence RTCA CardioECR System (Agilent). A 48-well CardioECR E-Plate was coated with 500 ug fibronectin from bovine plasma (Sigma). iCMs from an individual with PGM1 and a control were seeded 50,000 cells/well and cultured in RPMI-1640 media supplemented with B27 supplement, antibiotic-antimycotic, and 50 ug/mL uridine. After 7 days, well impedance was measured 1 ms-1 for 20 seconds.

### iCMs lysis and protein digestion

PGM1-deficient and healthy control iCM were first solubilized in 8 M urea (in 100 mM TEAB buffer) supplemented with 1% protease inhibitor cocktail (Thermo Scientific) and then sonicated with a tip sonicator at 30% amplitude for 3 cycles of 10 seconds each. Cells were centrifuged at high speed to remove the cell debris. Protein amount was estimated by BCA assay as per the manufacturer’s instructions (Thermo Scientific). Equal amounts of protein from both groups were first reduced with 10 mM dithiothreitol (Sigma-Aldrich, USA) at 37° C followed by alkylated with 40 mM iodoacetamide (Sigma-Aldrich, USA) at room temperature in dark. The proteins were then digested with 1:20 w/w (protein: trypsin) ratio of trypsin (Worthington, USA) at 37° C overnight. Resulting peptides were cleaned up using C18 cartridges and labeled with tandem mass tags (TMT) (Thermo Fisher Scientific, USA) as per the manufacturer’s protocol.

### Peptide fractionation

After checking the TMT labeling efficiency, the samples were subsequently pooled. Pooled peptides were subsequently split into two aliquots. One aliquot containing about 20% of total peptides was resuspended in solvent A (5 mM ammonium formate, pH 9) and fractionated by basic pH reversed phase liquid chromatography (bRPLC) on a C18 column (5 µm, 4.6 × 100 mm column, Waters) using a linear gradient of solvent B (5 mM ammonium formate, pH 9, in 90% acetonitrile) for 120 min on the Ultimate 3000 UHPLC system. Ninety-six fractions were collected and subsequently concatenated into 12 fractions. These concatenated 12 fractions were lyophilized and resuspended in 0.1% formic acid for liquid chromatography-tandem mass spectrometry (LC-MS/MS) for the proteomics study.

### Glycopeptide enrichment

The other aliquot of total peptides containing the remaining 80% peptides was resuspended in 0.1% formic acid and injected into Superdex peptide 10/300 column (GE Healthcare) as described previously[19,20,24]. The peptides were separated using an isocratic flow of 0.1% formic acid for 130 min and 12 early fractions were collected starting at 10 minutes after injection for N-glycoproteomics study. The fractions were lyophilized and resuspended in 0.1% formic acid for LC-MS/MS analysis for the N-proteomics study.

### Liquid chromatography tandem mass spectrometry (LC-MS/MS)

LC-MS/MS analysis of fractionated and enriched samples from both proteomics and N-glycoproteomics was carried out as previously described[19,20,24] with some modifications. Peptides were then separated by liquid chromatography on an EASY-Spray column (75 m × 50 cm, PepMap RSCL C18, Thermo Fisher Scientific) at a flow rate of 300 nl/min for 150 min using a linear gradient of 0.1% formic acid in water (solvent A) and 0.1% formic acid in acetonitrile (solvent B). The samples were analyzed on an Orbitrap Exploris 480 mass spectrometer (Thermo Fisher Scientific) equipped with Ultimate 3000 liquid chromatography system (Thermo Fisher Scientific Inc.). All experiments were done in DDA mode at an isolation window of 0.7 m/z. Precursor ions were acquired in the Orbitrap mass analyzer in m/z range of 300-1,700 for proteomics and 350-2,000 for N-glycoproteomics. Precursor ions were acquired at a resolution of 120,000 (at m/z 200) and fragment ions at a resolution of 30,000 (at m/z 200). Precursor fragmentation was carried out using normalized higher-energy collisional dissociation (HCD) method of 34 for proteomics and normalized stepped HCD at 15, 25 and 40% for glycoproteomics.

### Proteomics and glycoproteomics data analysis

Proteomics and N-glycoproteomics data analysis was performed as described previously[19,20,24]. The proteomics data were searched using Sequest search engine in Proteome Discoverer 3.0 against the human Uniprot protein database. The N-glycoproteomics data using the publicly available software pGlyco version 3 with an in-built N-glycan database for identifying glycans and human Uniprot protein database for identifying peptide sequence. Two missed cleavages were allowed for both proteomics and glycoproteomics analysis. Error tolerance for precursor and fragment ions were set to 10 ppm and 0.02 Da, respectively, for proteomics and 10 ppm and 20 ppm, respectively, for glycoproteomics. Cysteine carbamidomethylation was set as fixed modification, whereas oxidation of methionine as variable modification. False discovery rate (FDR) was set to 1% at the peptide-spectrum matches (PSMs), peptide, protein and glycopeptides levels. For proteomics, quantitation of peptides across PGM1-CDG and control fibroblasts was done using TMT reporter ion intensities using “reporter ion quantifier” node. To quantify glycopeptides, reporter ion quantification was performed for glycoproteomics raw files in Proteome Discoverer and glycopeptide IDs obtained from pGlyco were matched with quantitation on a scan-to-scan basis (MS/MS). For mitochondrial proteins, annotated gene list was generated from MitoCarta 3.0[25].

### MitoCarta analysis

To assess for a depletion of the MitoCarta[25] protein pool in our proteomics dataset, the number of MitoCarta protein species detected with a FC greater or less than 1 relative to controls was counted and compared to the number of total protein species detected displaying a FC greater than or less than 1 relative to controls. These counted proteins were then compared using Fisher’s exact test. This protocol was repeated for the detected subunits of mitochondrial electron transport chain complex I, complex II, complex III, complex IV, and complex V, and the mitochondrial ribosome, as well as all proteins involved with mitochondrial membrane integrity with an associated mitochondrial disease. These subunit proteins abundances and counts were compared to both the MitoCarta and total protein pools. Additionally the average FC relative to controls of all MitoCarta proteins, the detected subunits of mitochondrial electron transport chain complex I, complex II, complex III, complex IV, and complex V, and the mitochondrial ribosome, as well as all proteins involved with mitochondrial membrane integrity with an associated mitochondrial disease[26] was compared to the average FC of all protein species detected relative to controls using Kruskal-Wallis test.

### Ingenuity Pathway Analysis (IPA)

The functional analyses of the proteomics results were generated through the use of IPA[27] (QIAGEN Inc., https://www.qiagenbio-informatics.com/products/ingenuity-pathway-analysis). The IPA analysis was conducted using IPA default settings was performed on the top significantly changing proteins identified by proteomics. Log2 fold change of proteins with p-value of < 0.05 was used to calculate IPA analysis-ready molecules, and z-scores, representing the activation state of each canonical and disease pathway (z-score ≥ 2 significantly activated and z-score ≤ −2 significantly inhibited). All proteins with significant p-value <0.05 were analyzed. IPA analysis was limited to databases related to humans. To avoid bias, no tissue specificity was selected in the analysis. In addition, toxic (tox) functions analysis was performed by IPA, which catalogs the genes/molecules/proteins known to be involved with specific type of toxicity and their causal associations, when known. These analyses include associations to organ injuries, clinical chemistry and hematology assays, and pathway-related endpoints such as mitochondrial function etc. The molecules and their associations with different toxicity end-points and pathologies are hand-curated by IPA, and include renal, hepatic and cardiac injury, etc.

### GSEA analysis

The GSEA analyses of the proteomics results were generated using R (Version 4.3) an R-studio (V2023.09.1+494). For GSEA, a pre-ranked list based on log2foldchange*-log10(p-value) was used as input for the gseGO() function. Biological Process of GO Ontology was used as pathways. Benjamin Hochberg was used to calculate statistics. The adjusted p-value cutoff was set at 0.001 for upregulated pathways and 0.0001 for downregulated proteins. Semantically similar pathways were manually curated. A complete list of pathways can be found in Additional Table X. The following packages were used to create the graphs: ggplot2 (3.5.1), ClusterProfiler (4.10.1)[28].

### Tracer metabolomics experiments

Briefly, approximately 1million iCMs per well was plated on Geltrex coated 24-well plates in triplicates (day 0). The following day, the medium was refreshed (day 1). The medium was refreshed again after 48h (day 3). By day 5, all iCardiomyocyte cell lines were beating. The cells were washed with DPBS, and incubated with RPMI+insulin medium (ThermoFisher, 221704) supplemented with A) 5.5mM (physiologic concentration) ^13^C_6_-glucose, or B) 5.5mM ^12^C_6_-glucose. The cells were incubated for additional 48 h to ensure steady state labeling of the cells has occurred and metabolite extraction was performed as described below.

### Metabolite extraction

Briefly, cells were washed with ice-cold saline solution, 200 microL ice-cold extraction buffer (80 % MeOH, IS) was added each well. After 2 min, the cells were scraped in extraction buffer, transferred to fresh Eppendorf tubes and placed overnight at −80 °C. Next, all the samples were pelleted at 15,000 rpm for 20min at 4 °C. Supernatant containing metabolites was transferred to a new Eppendorf tube and metabolites measured with LC/MS (see below). 100 µL of 200mM NaOh was added to the cell pellets. The pellets were then incubated at 95 °C for 30min, centrifuged at 5000 rpm, rpm for 10min at 4 °C. Finally, the supernatant containing proteins was used for protein concentration determination with BCA Pierce Protein Assay kit (ThermoFisher, A55865).

### Metabolite measurement by LC/MS

Previously described method was used to analyze metabolites by LC/MS [20,29–31]. Briefly, 10 µL of sample was separated on a C18 ion-pairing liquid chromatography column and the metabolites resolved by Thermo Fisher Q-Exactive Hybrid Quadrupole Orbitrap MS in negative ion mode (resolution 140,000 at 200 m/z, AGC at 3e^6^, 512 ms ion fill time, full scan 70-850 m/z). The ESI settings were set to: 50 sheet gas flow rate, auxiliary gas flow rate 15, spray voltage of 4 kV, S-lens RF level of 60, and the capillary temperature at 350°C. Metabolite identification was performed according to their elution times, m/z ratio and in-house metabolite standard library. Peak picking and correction of naturally occurring carbon isotops was performed by El-Maven v0.12.0/Polly ^TM^ Labeled LC-MS Workflow[32]. Metabolite abundances were normalized to protein concentration and internal standards. Absolute quantification was not performed. Relative values were established using healthy control organoids as reference.

### Fractional contribution of ^13^C_6_-glucose and isotopologues statistical analysis

Fractional contribution of ^13^C_6_-glucose is defined as a percentage of ^13^C_6_-glucose contributing to the pool of specific measured metabolite. Isotopologue labeling (positional labeliing) of ^13^C_6_-glucose is defined as a percentage of numbers of carbons labeled by ^13^C_6_-glucose in a specific metabolite (m0-mn, where n signifies the number of carbons in a specific metabolite). Visualization of the fractional contribution (FC) of ^13^C_6_-glucose in PGM1-CDG and control iCMs, was performed using TraVis Pies[33]. TWO-way ANOVA was used to assess the significant differences between PGM1-deficient and CTR iCMs in overall isotopologue labeling and significant difference in the isotopologues distribution based on the genotype (interaction between genotype and isotologue distribution). Multiple comparisons with Sidak correction analysis was used to assess significant differences between PGM1 deficient and CTR iCMs in specific isotopologues. The results of the statistical analysis are provided in additional data.

### Metabolic pathway analysis

MetaboAnalyst 5.0[34] was used to perform pathway analysis on metabolomics data. Metabolomics data containing FC information and metabolite IDs was analyzed by standard MetaboAnalyst parameters. These included: enrichment method-global test; topology measure relative-betweenness centrality, and the metabolites were compared to all the compounds in the KEGG pathway library (Homo Sapiens).

### Mitochondrial respiration studies

Seahorse XFe96 Extracellular Flux Analyzer (Agilent, Santa Clara, CA, USA) and XF Cell Mito Stress Test kit (Agilent, 103015-100) were used to investigate the mitochondrial respiration of PGM1-CDG and control iCMs, as previously described in patient fibroblasts[35]. The seeding density and inhibitor concentrations were specifically optimized for iCMs. Briefly, both control and PGM1 iCMs were simultaneously thawed as described above, and seeded at 40,000 cells per well in RPMI+ insulin supplemented with 10% KOSR and 10µM ROCK inhibitor in a 96-well Seahorse microplate (day 0). The medium was refreshed the following day (day 1), and once again after 48h (day 3). Then, 48h before the measurements, the medium was changed to RPMI+ insulin containing 5.5 mM (physiologic concentration) glucose (day 5). On the day of the measurement (day 7), the cells were washed with DPBS and the medium was replaced with XF Base Medium Minimal RPMI (Agilent, 103576-100) supplemented with 10 mM XF seahorse glucose (Agilent, 103577-100), 1 mM XF seahorse pyruvate (Agilent, 103578-100), and 2 mM L-glutamine according to the manufacturer’s instructions. After the initial basal respiration measurements, the specific inhibitors used to assess mitochondrial respiration were added in following order: port A 2.5 μM oligomycin, port B 2.0 μM carbonyl cyanide phenylhydrazone (FCCP), port C 0.5 μM rotenone + antimycin A (Agilent, 103015-100). After the measurements, the cell membrane-permeable nuclear staining compound (Hoechst 33342) was added to the cells and cell counting using Celigo imaging cytometer (Beckman) was performed. Each cell line was seeded in 8-well replicates. The experiment was repeated 3 times using new cells each time, avoiding freeze-thaw cycles. The results were normalized to the cell number and citrate synthase (CS) activity, a proxy readout for mitochondrial mass[36], described below.

### Citrate Synthase (CS) activity/abundance

CS is a matrix enzyme of the TCA/Krebs cycle that catalyzes the conversion of oxaloacetate and acetyl-CoA to citrate and CoA. CS is highly enriched in the mitochondrial matrix and is thus commonly used as a proxy readout for mitochondrial mass[36]. Cell lysates were incubated with acetyl-CoA and oxaloacetate in the presence of DTNB (5’,5’dithiobis-(2nitrobenzoaat)).Precipitation of acetyl-CoA and oxaloacetic acid to citrate releases S-CoA, which cleaves DTNB (colorless) to TNB^2-^ (yellow). Increase in absorption at 412 nm was assessed as CS activity. For a blank measurement, the reaction was performed in the absence of oxaloacetate.

### *In silico* prediction of LDB3-PGM1 interaction using AlphaFold3

AlphaFold3 (AF3)[37] was used to predict potential interactions between PGM1 and Zasp/Cypher (LBD3). In particular, we focused on potential interactions with Exon 4 of Zasp/Cypher, which have been previously identified[17]. Exon 4 contains the conserved ZASP motif (residues 189-214 of LBD3 isoform 1), implicated in protein interactions. AF3 used on PGM1 predicts structure with high confidence, as expected give that its crystal structure is known. AF3 used on Exon4 (of unknown structure) indicates mostly disordered regions but does predict a structured region with high confidence that corresponds approximately to the ZASP motif. In predictions of a potential PGM1-Exon 4 complex, the top scoring model returned values of 0.45 for ipTM (just below the 0.5 level suggesting a correct prediction) and pTM of 0.80 (matching the 0.8 cutoff for a confident high-quality prediction) using the PGM1 sequence (isoform 1). Moreover, the predicted aligned error (PAE) for the interaction between PGM1 and the ZASP motif of Exon 4 is low (<5 Å) supporting an accurate prediction (Fig. 2D). (As noted in AF3 documentation, PAE for may be more reliable for complexes involving small partners.) We also tested the entire LBD3 sequence as well as Exon 10 in the AF3 server but did not find confident predictions of a complex with PGM1.

**Figure 2.**
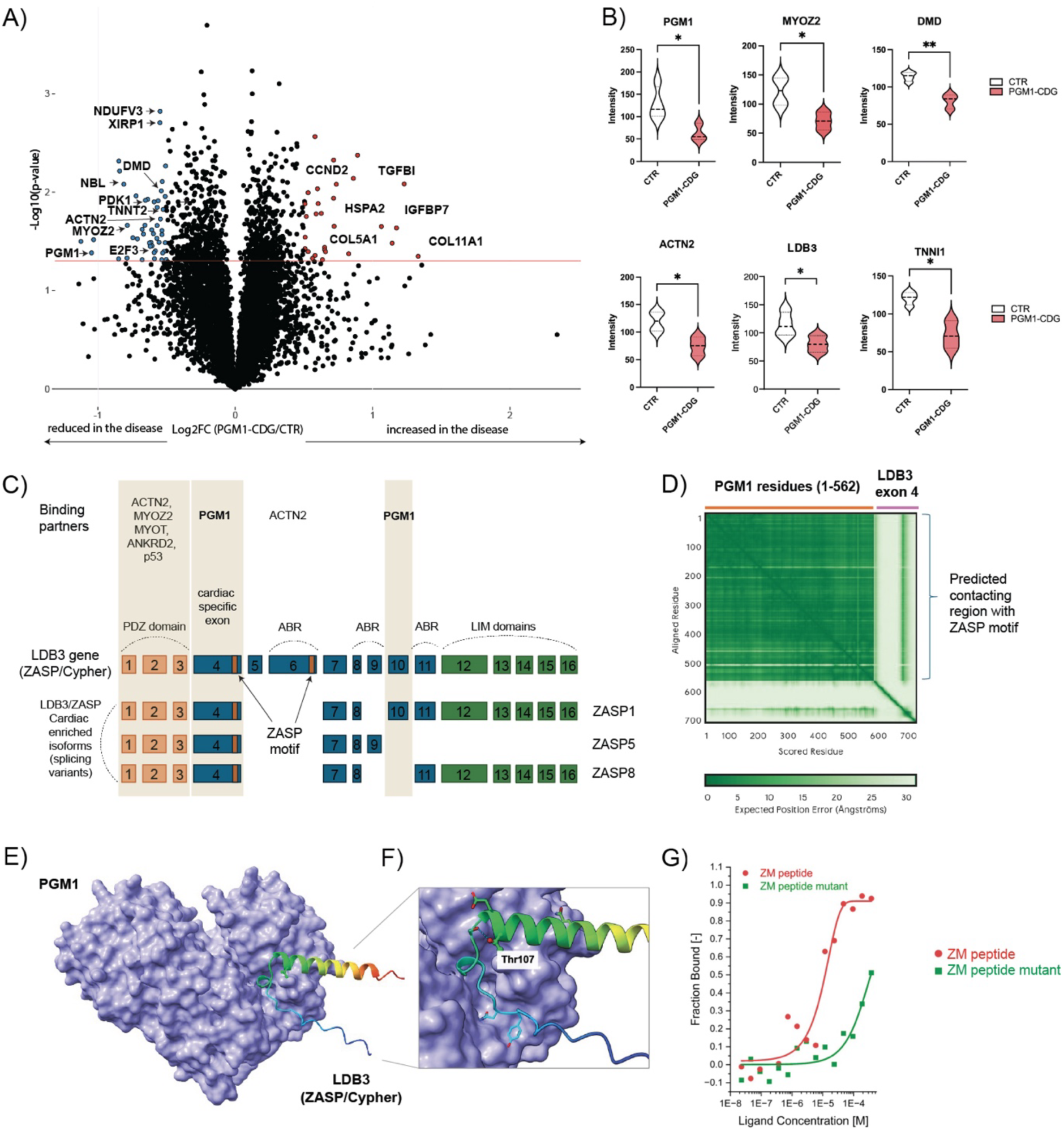
Sarcomeric Protein Dysregulation and Potential Z-Disk Impairment in PGM1-Deficient iCMs. **A)** Volcano plot showing differentially abundant proteins in PGM1-deficient iCM compared to healthy control iCM (CTR). X-axis shows log_2_ fold change (FC) (PGM1-deficient/CTR) and Y-axis represents the −log10 of p-value. The horizontal red line represents the cutoff for significance [-log10(0.05)]. The most significantly changing proteins are marked in either red (>1.3 FC, p< 0.05) or blue (<0.7, p<0.05). **B)** Violin plots showing some of the top significantly changing proteins involved in cardiomyocyte machinery in PGM1-deficient iCM compared to CTR. Y-axis represents the ion intensity of TMT channels. PGM1 n=3, t=1; CTR n=4, t=1. Violin plots are shown with standard deviation (SD). Detailed statistical analysis can be found in the supplemental information. **C)** Schematic of the LDB3 gene (16 exons) that encodes various isoforms of ZASP/Cypher. The three splicing variants of ZASP (isoforms) are also shown with the cardiac specific exon 4 and ZASP motifs on exon 4 and 6 indicated by orange box. Regions with known interactions with PGM1 (exons 4 and 10), as well as ACTN2, MYOZ2, MYOT, ANKRD2 and p53 are highlighted. ABR-actin binding region. **D**) The predicted alignment error (PAE) matrix from AF3 for the top scoring model of the PGM1 and Exon 4 complex, which had ipTM and pPTM values of 0.45 and 0.80, respectively. **E**) A figure showing the predicted interaction between PGM1 and the ZASP motif of Exon 4 of LBD3 (Zasp/Cypher). PGM1 is shown in a surface representation (purple); residues 85-128 of Exon 4 are shown as a ribbon, in a color ramp from blue (N-terminus) to red (C-terminus). The ZASP motif is found within the helical region (green) and Thr107 is shown in a space filling model. A Thr to Ile mutant of residue 107 has been reported to impair interactions with PGM1[17]. **F**) A close-up view of the potential PGM1 interaction with the ZASP motif of Exon 4. The location of Thr107 near the N-terminus of the helix is highlighted, along with a hydrogen bond between the Thr107 side chain and a preceding residue. This interaction suggests that Thr107 may play a role in stabilizing the helical conformation of the ZASP motif and that mutation to Ile could disrupt the helical structure and impair binding to PGM1. G) High affinity *in vitro* interactions between exon 4 of LBD3 (ZASP/Cypher) and PGM1.

### Microscale-thermophoresis

For microscale-thermophoresis studies, a peptide representing the conserved Zasp motif (ZM) of ZASP/Cypher (26-mer: SSQPRQYNNPIGLYSAE<u>T</u>LREMAQMY) and a corresponding peptide with the T106I mutation (underscored in sequence) were synthesized by ABI Scientific. A vector for recombinant expression of human PGM1 has been previously described[38]. PGM1 was expressed and purified as previously described[38].

For binding studies, PGM1 was labeled using a commercial kit (NanoTemper Technologies, Inc.) according to the manufacturer’s specifications. Briefly, PGM1 was incubated at room temperature for 30 minutes with RED-tris-NTA dye, which specifically labels His_6_ tags, resulting in 100 nM labeled protein. The wild-type and T106I ZM peptides were serially diluted into a series of 16 PCR strip tubes and mixed 1:1 with labeled PGM1. The final concentration series of each peptide was 375 mM to 11.4 nM, with all tubes containing 50 nM labeled PGM1. Samples were incubated in the dark for 10 minutes prior to use. Standard capillaries (Nanotemper MO-K022) were used to draw up the liquid and placed onto the sample tray of the Monolith NT.115 instrument (NanoTemper). Measurements were performed at 22°C with control software at 40% excitation power and 40% MST power. K_d_ values were calculated using the MO.Affinity software v.2.2.4 (NanoTemper Technologies).

### EMUDRA in silico drug repurposing

Ensemble of Multiple Drug Repositioning Approaches (EMUDRA) [39] algorithm was applied to the proteomics data using R (Version 4.3) an R-studio (V2023.09.1+494). For EMUDRA Fig 7A, the significantly overexpressed proteins were used as input (log2foldchange >0.25 and p-value 0.05). For EMUDRA Fig 7B, the significantly decreased proteins were used as input (log2foldchange <0.25 and p-value 0.05). The following packages were used to create the graphs: ggplot2 (3.5.1), EMUDRA (1.0.5) and EMUDRA data (1.5).

### Statistical analysis and visualization

GraphPad prism 10 for MacBook was used for the statistical analysis and violin plot generation, unless otherwise specified. Unpaired t-test with unequal group variance was used to analyse proteomics and glycopeptide data (FDR <0.05). Mitocarta proteins were also analyzed by Fisher’s exact test and Kruskal-Wallis (multiple t-test Dunn’s multiple comparisons test). For Seahorse oxymetry, Shapiro-Wilk test was used to determine population normality/lognormality, and multiple unpaired either parametric or non-parametric T-tests were performed with a false discovery rate of 1%. Two-way ANOVA was performed to analyze sources of ATP. For tracer metabolomics data, unpaired t-test with unequal group variance was performed for the metabolite abundance and fractional contribution. TWO-way ANOVA with multiple t-test and Sidak correction was performed for positional labeling. Further [33]pathway analysis was performed with Metaboanalyst[34]. Volcano plots were generated in R, using ggplot2 package. Means are represented with SD. Biorender with licence was used to generate parts of the figures. Figures were further prepared using GraphPad, MetaboAnalyst, Travis Pies[33] and Adobe Illustrator. The details of statistical analysis for each experiment can be found in Additional Spreadsheet.

## Results

### Clinical presentation of PGM1-CDG patients included in the study

The fibroblasts of four previously reported unrelated individuals affected with PGM1-CDG were used to generate iCMs for this study. The demographic and biochemical data of the four individuals with PGM1-CDG are listed in **Table 1**. All four individuals had pathogenic variants in *PGM1* affecting both PGM1 isoforms (PGM1-1, PGM1-2)[10]. The oldest patients reported were 49[40] and 53-years-old [8] at the time of the first symptoms. Therefore, yearly monitoring of cardiac function has been recommended in all patients affected by PGM1-CDG[1].

**Table 1.** Characteristics of the four individuals with PGM1-CDG whose fibroblasts were used to generate iCMs. *age at the time of the first publication, #variants reported in the first publication

| ID | Age*/Gender | PGM1 variants<sup>#</sup> | PGM1 amino acid change | Cardiac presentation |
| --- | --- | --- | --- | --- |
| P1[5,14,41] | 3/F | c.551delT<br>HOM | p.Phe184Serfs*9 | No cardiomyopathy at the time of assessment |
| P2[5,14,41] | 10/M | c.1162G>A<br>c.1547T>C | p.Glu388Lys<br>p.Leu516Pro | CM-severe<br>FS <10%. Cardiac arrest- deceased |
| P3[5] | 16/F | c.1508G>A<br>HOM | p.Arg503Gln | Left ventricular hypertrophy |
| P4[5] | 20/F | c.1145-222G>T<br>HOM | p.Gly382ValfsTer23 | Severe cardiomyopathy<br>FS 7%, cardiac arrest- resuscitated |

In our cohort, three out of four patients presented with cardiac involvement. (P2-4). The youngest individual (P1), a three-year-old female, did not present yet with cardiac involvement at the time of PGM1-CDG diagnosis. However, based on her molecular findings and her young age, this individual is likely to develop cardiomyopathy[10]. P2 had severe cardiomyopathy (ejection fraction <10%). He suffered a cardiac arrest and was deceased at the age of 10. P3, a 16-year-old female, presented with left ventricular hypertrophy. P4 is a 20-year-old female who also presented with severe cardiomyopathy (fractional shortening 7%) and suffered from cardiac arrest but was resuscitated.

### Differentiation of PGM1-deficient hIPSC into iCardiomyocytes (iCMs)

The successful generation hIPSC from the fibroblasts of four individuals with PGM1-CDG was confirmed by quality control tests including karyotyping and mycoplasma testing (see methods for details). The pluripotency markers and the ability of the hIPSC to differentiate into three germ layers were assessed (**Fig 1B)**. The PGM1-CDG (n=4) and control (CTR, n=4) hIPSC were differentiated into iCMs by using previously reported chemically defined protocol[21,22] (see methods for details). The schematic diagram demonstrating the main steps of the protocol used for the direct differentiation of hiPSC into to iCM is shown in **Fig 1C**. Successful differentiation of hiPSC into iCM was confirmed by assessing cardiomyocyte markers[42] by gene expression analysis, myosin light chain 6, myosin heavy chain 6, myosin heavy chain beta, and troponin) (**Fig 1D**) and immunohistochemistry (connexin-43, alpha-actinin, cardiac troponin) **(Fig 1E**).

### PGM1-deficient iCM exhibit decreased beating frequency and contractility

Like cardiomyocytes, iCMs generate action potentials, resembling heartbeats in the dish, which can usually be observed by naked eye. However, more advanced methods such as multi-electrode assay analysis (MEA) are used to quantify the action potential of iCMs in the dish, which is specifically useful in the case of e.g., drug screening [22,43,44].. While generating the PGM1-deficient iCM, we noticed that PGM1-deficient iCM had a lower yield of iCMs and started beating later than the CTR iCMs or would not beat at all. To assess the functional consequences of PGM1 deficiency, we performed MEA in both CTR and PGM1-deficient iCMs (**Fig 1F**). We found that PGM1-deficient iCM had a significantly lower ‘heartbeat’ count compared to the healthy controls (**Fig 1G**), confirming that PGM1-deficient iCMs have a profound cardiac dysfunction, like the one reported in the PGM1-CDG individuals[1]. Representative CTR and PGM1-deficient MEA recordings are shown in (**Fig 1H**). In addition, PGM1 iCMs displayed a profound reduction in contractility relative to the control (**Fig 1I).** Stimulation with 0.3 ms pulses of 1200 mV at either 0.5 or 1.0 Hz had no effect on contraction amplitude (**Fig 1J**). PGM1 iCMs displayed increased time between application of the stimulation and both contraction initiation and peak contraction. PGM1-iCMs also displayed increased time between peak contraction and both 10% and 50% relaxation (**Fig 1K**). These data suggest that PGM1-iCMs display a slower and less powerful contraction, consistent with a bradycardic phenotype and cardiac dysfunction.

### Sarcomeric Protein Dysregulation and Potential Z-Disk Impairment in PGM1-Deficient iCMs

As PGM1 iCM already displayed severe cardiac dysfunction during the differentiation, we sought to investigate the consequences of PGM1 deficiency on a molecular level. To assess the global effect of PGM1 deficiency, we first performed a multiplexed proteomic analysis in PGM1-deficient and healthy CTR iCMs. We identified 7751 unique proteins across all samples and found profound changes in global protein expression in PGM1-deficient iCMs (**Fig 2A**, **Sup Fig 1**). PGM1 was among the top downregulated proteins (average FC=0.48) in all PGM1-deficient iCMs as compared to CTR (**Fig 2B)**. Consistent with cardiac dysfunction observed with MEA, several proteins involved in cardiomyocyte function and sarcomere formation were significantly decreased in PGM1-deficient iCMs such as DMD, XIRP1, TNN1, TNNI1, TNT2, ACTN2, MYOZ2 (**Fig 2A, B**). Apart from proteins involved in cardiomyocyte architecture, several of the top significantly increased proteins (FC >1.3, p<0.05) in PGM1-deficient iCMs belonged to the proteins involved in extracellular matrix (ECM) formation or collagen synthesis such as IGFBP7, TGFBI, COL5A1 and COL11A1. On the other hand, cell cycle and cell cycle progression including E2F3, CEP290, SASS6, Aurora and cyclin D2 (CCND2) (**Fig 2A**) as well as proteins involved in mitochondrial function and signaling (e.g. NDUFV3, SDHAF4, MCT2, ACAD10, COX4I1) were decreased in PGM1-deficient iCMs (**Fig 2A, Sup Fig 1**).

Previously, the cardiac specific PGM1 KO model showed Z-disk disarray and swollen/fragmented mitochondria on histology[19]. Given that PGM1 was previously linked to sarcomere Z-disk protein LDB3 (ZASP/Cypher)[17], we were specifically interested in further assessing the possible relationship of PGM1 with ZASP/Cypher. Previous studies showed PGM1 colocalizes with LDB3 on immunohistochemistry in the Z-disk[17]. Pathogenic variants in *LDB3* have been shown to disrupt the interaction of PGM1 and LDB3, and result in sarcomere disruption and dilated cardiomyopathy[17]. In PGM1-deficient iCMs, LDB3 was decreased (average FC=0.69) (**Fig 2B**). In addition, ACTN2 and MYOZ2[18], both known to interact with LDB3 in the Z-disk (**Fig 2 B**) were decreased suggesting Z-disk might be compromised in PGM1-deficient iCMs. To further assess the interaction between PGM1 and LDB3, and link PGM1 deficiency to sarcomere disruption and cardiac dysfunction, we also performed *in silico* interaction analysis using AlphaFold3[37]. ZASP protein has several isoforms, including cardiac-specific isoforms, and a cardiac-specific exon 4 (Zasp motief ZM), which is thought to bind PGM1 (**Fig 2C).** AlphaFold3 identified a potential binding site of PGM1 to LDB3 cardiac-specific exon 4 **(Fig 2 D, E, F**), which was further confirmed by *in vitro* binding assay experiments with PGM1, LDB3 and mutant LDB3 (**Fig 2G**). LDB3 (Zasp-motief ZM) peptide from exon 4 (red) showed high affinity (low μM) binding to PGM1. In addition, the peptide with the DCM-related *LDB3* T206I mutation (ZM mutant) showed decreased binding affinity (green) (**Fig 2 G**). These are the first *in vitro* data on interactions between PGM1 and ZASP/Cypher, confirming their proposed binding and the deleterious effect of a *LDB3* cardiomyopathy-related mutant.

Together with previous reports[17], this data suggests PGM1 could be directly involved in Z-disk function through binding with sarcomere Zasp protein LDB3. As PGM1 binds to a cardiac-specific region of LDB3, these findings also partially explain the cardiac-specific consequences of PGM1 deficiency in the heart, independent of glycosylation abnormalities.

### PGM1-deficient iCMs present with mitochondrial dysfunction

Sarcomere is a complex structure consisting of multiple proteins. Sarcomere contraction requires a lot of energy and the close proximity of mitochondria to the sarcomeres ensures that energy is readily available. Recently, the importance of mitochondria in sarcomere organization and function was also suggested [45]. Considering the importance of mitochondria and their close proximity to sarcomere[46], it is not surprising that the cardiomyocyte architecture disruption has been linked to mitochondrial dysfunction[47,48]. Particularly, pathogenic variants in sarcomeric genes, including LDB3, were shown to also result in mitochondrial abnormalities [49–51]. Analogously, mitochondrial abnormalities have previously been shown using histology in a heart of a PGM1-CDG patient, who underwent transplantation[19], as well as in a cardiac-specific PGM1 KO mouse [19].

To understand the extent of the mito-proteome changes in PGM1-deficient iCMs, we compared the proteomics findings with the MitoCarta protein pool, a catalogue of over 1000 genes encoding the mammalian mitochondrial proteome[25] (**Fig 3A**) (see methods for details). PGM1-CDG iCM’s displayed a profound depletion in the MitoCarta protein pool, with 84% of the 912 MitoCarta proteins identified displaying a fold change <1, compared to the 47% of the total protein pool displaying a fold change <1 relative to controls (**Fig 3A**), and an average depletion of 13% relative to the total protein pool (**Fig 3J**). Consistent with this depletion of the MitoCarta protein pool, we also observed significant depletions of the subunits of CI, CII, CIII, CIV, CV, and the mitochondrial ribosome (**Fig3 B-G**), with average depletions of 22%, 22%, 20%, 17%, 24%, and 18%, respectively (**Fig 3J**). Additionally, the subunits of CI and the mitochondrial ribosome displayed even further depletion relative to the already depleted MitoCarta protein pool (**Fig 3 B,G**), with average depletions of 11% and 6% relative to the MitoCarta protein pool respectively (**Fig 3J**).

**Figure 3.**
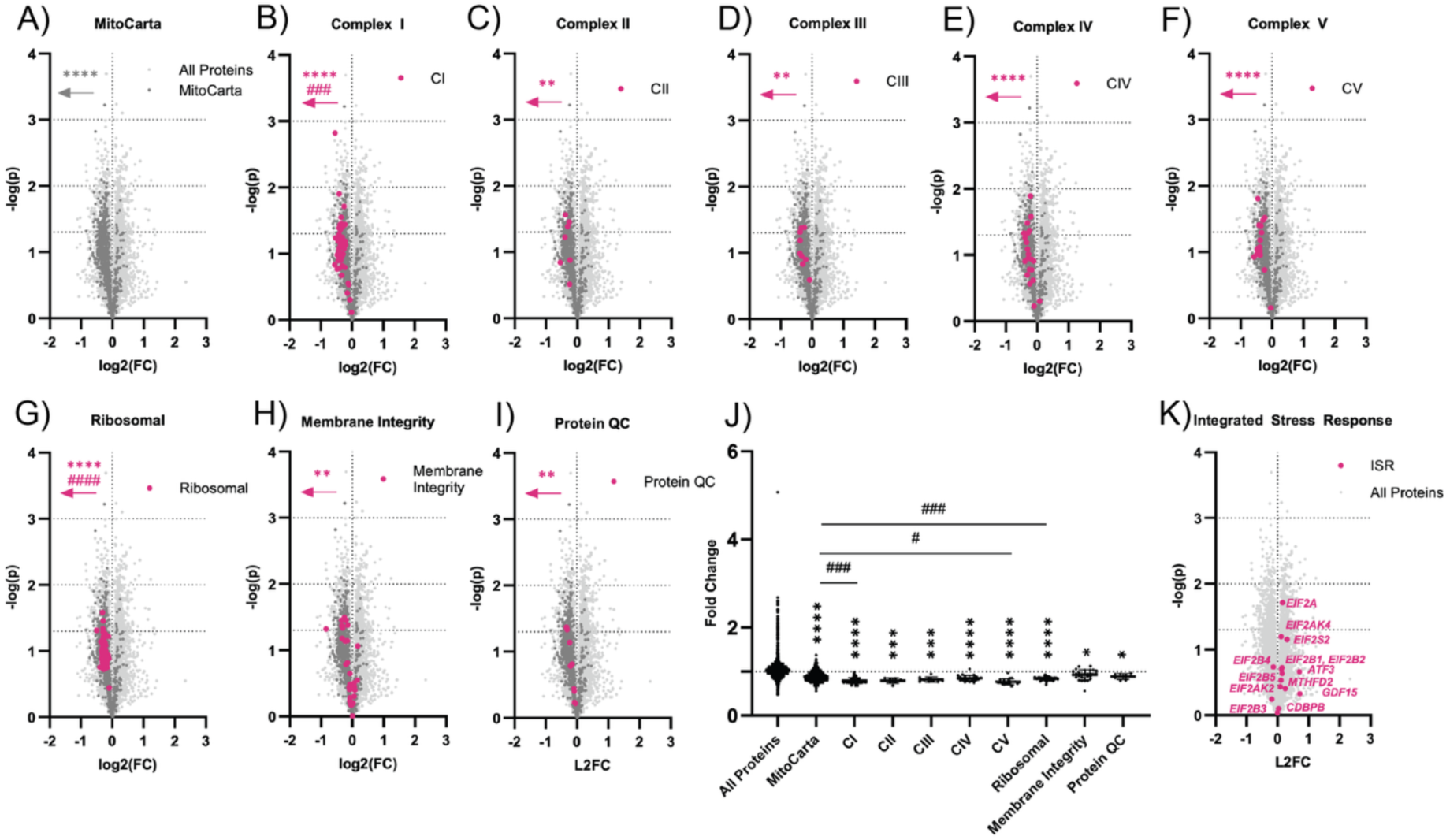
Mitochondrial proteins are depleted in PGM1-deficient iCMs. [17]A-I) Volcano plots displaying log2 fold change (FC) vs. −log p value (p) of all proteins (A), MitoCarta proteins (A), and the subunits of complex I (B), complex II (C), complex III (D), complex IV(E), complex V(F), and the mitochondrial ribosome (G) and all proteins associated with mitochondrial membrane integrity (H) and protein quality control (QC) (I) in PGM1-CDG cardiomyocytes (iCMs) relative to controls. Arrows indicate an increased number of depleted proteins species (Fisher’s exact test) of each subunit pool relative to the total protein pool (*) or the MitoCarta protein pool (#). J) Scatter plot of fold changes of all proteins, MitoCarta proteins, and the subunits of CI, CII, CIII, CIV, CV, and the mitochondrial ribosome, as well as proteins involved in mitochondrial membrane integrity and protein quality control in PGM1-CDG iCMs relative to controls. Significantly reduced average fold change (Kruskal-Wallis test) relative to all proteins (*) and the MitoCarta protein pool (#) are indicated. K) Volcano plots displaying log2(FC) vs. −log(p) of integrated stress response proteins in PGM1-CDG iCMs relative to controls.(#/*) p<0.05, (##/**) p<0.01, (###/***) p<0.001, (####/****) p<0.0001.

As mitochondrial structural abnormalities were found in the PGM1-CDG patient’s heart and cardiac-specific PGM1 KO mouse model[19], we assessed the expression of the proteins involved in mitochondrial membrane integrity, biogenesis/remodeling, protein import and protein quality control, as well as proteins involved in mitochondrial dynamics/interorganelle communication[26] (**Fig 3B**). Proteins associated with mitochondrial membrane integrity displayed significant depletions relative to the total protein pool (Fig2 H), with an average depletion of 9% (**Fig 3H**). Within this subset of membrane integrity-associated proteins, proteins associated with protein quality control (QC) were also depleted (**Fig 3I**), with an average depletion of 13% relative to the total protein pool (**Fig 3J**).

These results further corroborate the previously reported mitochondrial structural abnormalities in the PGM1-deficient heart[19].

Given the profound depletion in the total MitoCarta protein pool (**Fig 3B**), we hypothesized that the integrated stress response (ISR) pathway may be activated in PGM1-CDG iCMs. The ISR is activated in response to cell stress, including metabolic cell stress[52]. Activation of the ISR results in phosphorylation and subsequent sequestration of Eukaryotic Transcription Initiation Factor 2 Subunit Alpha (eIF2a), which is required for initiation of protein translation, reducing global protein translation. Activation of the ISR also induces the expression of multiple genes involved in alleviating cell stress, including AT3, ATF4, ATF 5, GDF15, FGF21, MTHFD2, and CDBPB. Relatively few components of the integrated stress response pathway were identified through proteomic analysis, including eIF2a, multiple eIF2 kinases, several subunits of eIF2B, as well as multiple downstream protein species, viz. ATF3, GDF15, MTHFD2, and CDBPB. Of these, only eIF2a displayed significant enrichment (12%, p=0.019) (**Figure 3K**). While the phosphorylation status of this enriched eIF2A could not be assessed, the absence of significantly enriched downstream proteins of the ISR pathway, i.e. ATF4, ATF3, ATF5, GDF15, FGF21, MTHFD2, and CDBPB, there is no evidence to suggest that the ISR pathway is induced in PGM1-CDG iCMs.

Together with previously reported data from cardiac specific PGM1 KO model and PGM1 patient heart[19], these results indicate that similarly to LDB3-related cardiomyopathy[49–51], PGM1-deficiency results in severe mitochondrial changes.

### Pathways analysis confirms mitochondrial dysfunction as the major hallmarks in PGM1-deficient iCMs

To avoid bias in assuming that the disruption of cardiac cytoskeleton, including sarcomere and mitochondria, were the major hallmarks of PGM1-deficient heart, we employed Ingenuity Pathway Analysis (IPA, QIAGEN, see methods) to independently assess the extent of the pathways affected in PGM1-deficient iCMs (**Fig 3A, B**). IPA analysis showed several canonical pathways were significantly changed in PGM1-deficient iCMs (**Fig 3A**). The top downregulated pathways were associated with oxidative phosphorylation, electron transport chain, ATP production and heat production by uncoupling proteins (**Fig 4A**). The top upregulated canonical pathways in PGM1-deficient iCMs were mitochondrial dysfunction, eukaryotic translation elongation, intra-GA and retrograde-GA to ER trafficking as well as protein sorting (**Fig 4A**). In addition, other pathways associated with cellular trafficking were significantly affected (**Fig 4A**).

**Figure 4.**
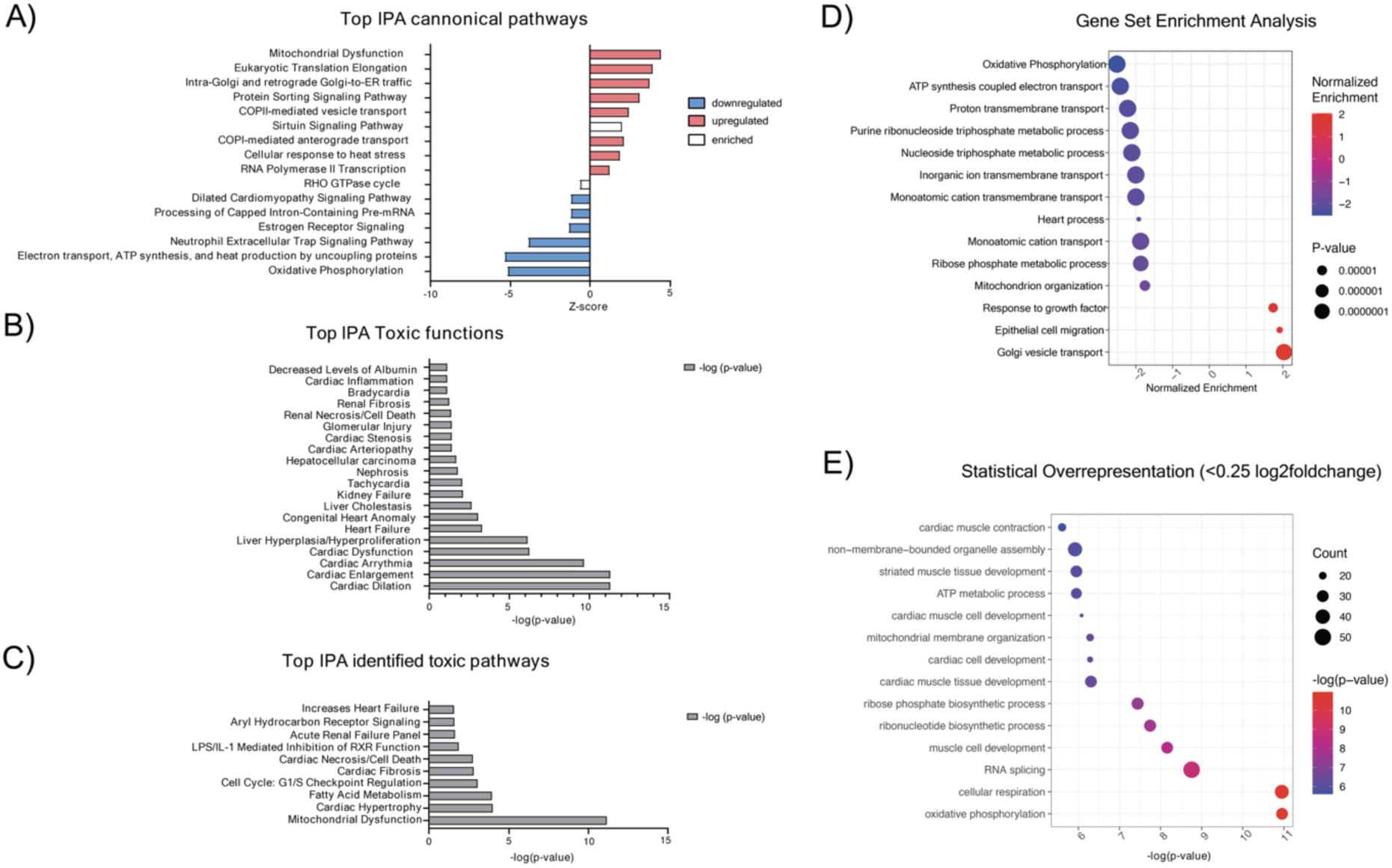
Pathway analysis of the significant proteomic changes in PGM1-deficient iCM. A) Top changing canonical pathways according to Ingenuity Pathway Analysis (IPA) The pink color represents upregulated pathways, while blue color signifies downregulated pathways. White color represents pathways that are enriched in the dataset, but the direction of their activity (upregulated/downregulated) cannot be predicted. B) IPA identified top significant toxic functions associated with the proteomic signature of PGM1-deficient iCM. C) IPA identified top significant toxic pathways based on the significant proteomics changes in the PGM1-deficient iCM D) GSEA results based on pre-ranked proteome log2foldchange- and p-values of proteomics data of PGM1-deficient iCMs compared to healthy controls. As pathways, all Biological Process (BP) pathways of the GO ontology were included. The graphs show pathways with p-value <0.0001 for downregulated pathways (corrected using Benjamin Hochberg method) and p-values <0.001 for upregulated pathways. (PGM1 n=3, t=1; CTR n=4, t=1)

Corresponding to the observed decrease of proteins involved in cardiomyocyte/sarcomere machinery (**Fig 2**), the top 5 upregulated toxic functions in PGM1-deficient iCMs were associated with cardiac dilation, cardiac enlargement, cardiac arrythmia, cardiac dysfunction and heart failure (**Fig 4B).** We also used IPA to search for the toxic (disease-causing) pathways that overlap with the signature of PGM1-deficient iCMs. The top toxic pathways associated with PGM1-deficient proteome were mitochondrial dysfunction and cardiac hypertrophy (**Fig 4C**).

In addition, we performed gene set enrichment analysis (GSEA)[53]. GSEA showed several significantly enriched pathways (**Fig 4D, E, F**). Downregulated pathways included oxidative phosphorylation, purine/nucleotide, ribose phosphate metabolism and inorganic ion transport. Upregulated pathways included response to TGF-beta, collagen and ECM pathways (**Figure 4E, F**).

Together, the IPA and GSEA analysis data suggested mitochondrial function, ribonucleotide metabolism and ECM organization are the most affected pathways in PGM1-deficient cardiomyocytes and therefore potential contributors of the severe cardiac phenotype observed in PGM1-CDG.

### PGM1-deficient iCMs show abnormal glycosylation

Our proteomics studies revealed profound depletion of proteins involved in cardiomyocyte architecture and mitochondrial function, which have previously been linked to cardiac dysfunction[50,54–60]. However, as PGM1 is directly involved in glycosylation and PGM1 deficiency results in glycosylation abnormalities treatable by galactose[11] **(Fig 5A**), we also sought to explore the effect of PGM1 deficiency on the glycosylation in iCMs. To do this, we performed untargeted glycoproteomic analysis in PGM1-deficient and CTR iCMs.

**Figure 5.**
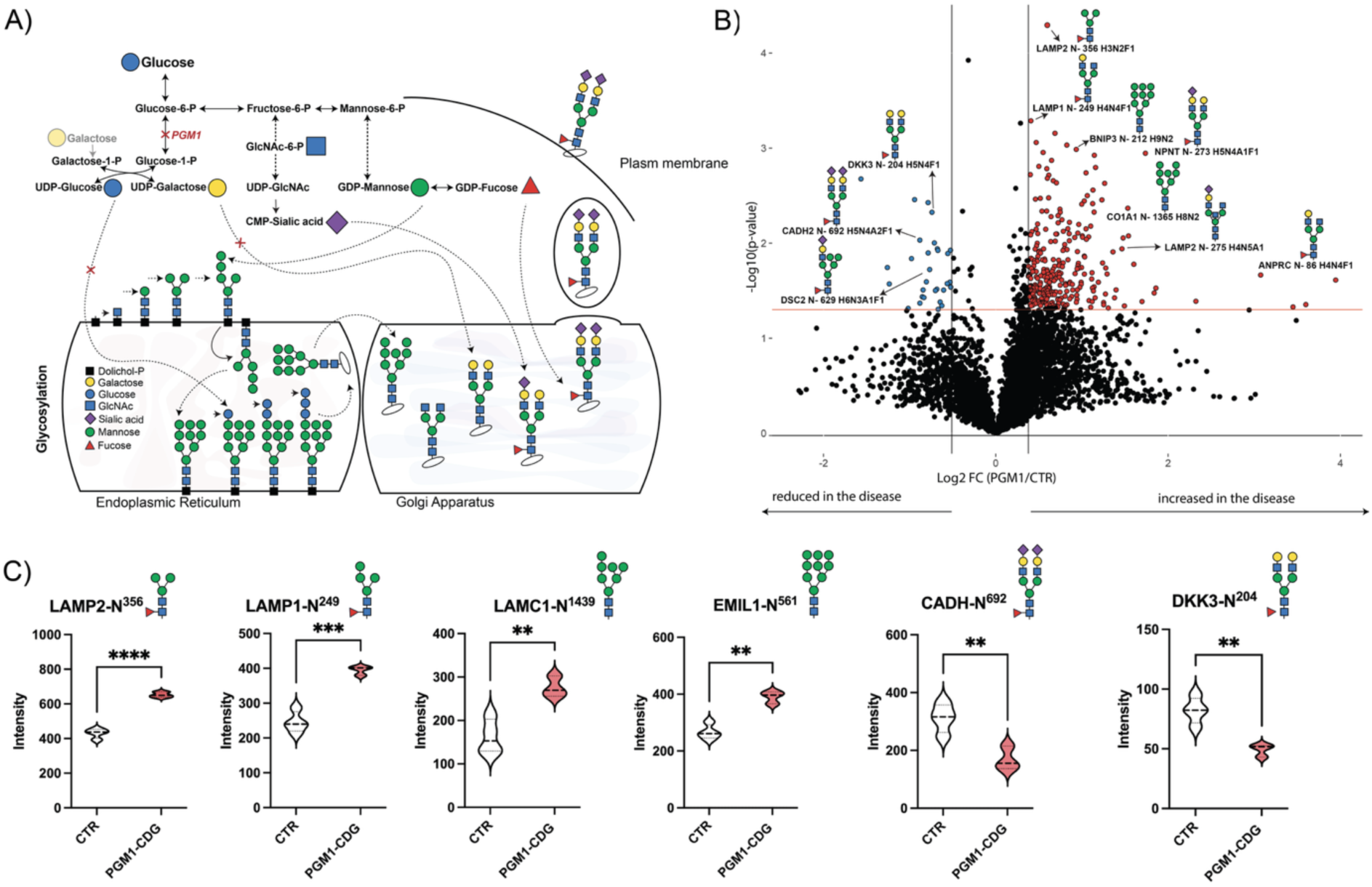
Global glycoproteomic changes in PGM1-deficient iCMs (iCM). A) Schematic representation of PGM1 in relation to glycosylation. Glycosylation starts in cytosol, where suger nucleotides (UDP-Glucose, UDP-Galactose, UDP-GlcNAc, GDP-Mannose, GDP-Fucose, CMP-sialic acid etc.) are produced. The sugar nucleotides serve as donors of sugars to the growing glycan chains in Endoplasmic reticulum (ER), where glycans are attached to proteins and then further processed in Golgi apparatus (GA). Fully formed glycans are exported to the cell surface. B) Volcano plot showing differentially abundant glycoproteins in PGM1-deficient iCM compared to healthy control iCM (CTR) after correction for the proteins that were significantly changing on protein level. X-axis shows log_2_ fold change (FC) (PGM1-deficient/CTR) and Y-axis represents the −log10 of p-value. The horizontal red line represents the cutoff for significance [-log10(0.05)]. The most significantly changing proteins are marked in either red (>1.3 FC, p< 0.05) or blue (<0.7, p<0.05). C) Violin plots showing some of the top significantly changing glycoproteins in PGM1-deficient iCM compared to CTR. Y-axis represents the ion intensity of TMT channels. PGM1 n=3, t=1; CTR n=4, t=1. Violin plots are shown with standard deviation (SD). Detailed statistical analysis can be found in the supplemental information.

We identified 4772 unique glycopeptides derived from 574 proteins across all samples. We found profound changes in glycosylation in PGM1-deficient iCMs compared to CTR, out of which the abundances of 486 unique glycopeptides were significantly changing (p<0.05) (Additional Data). As we identified profound changes in the proteome of PGM1-deficient iCMs, we then performed additional analysis, by filtering out the glycopeptides coming from proteins whose abundance was also significantly changed at the protein level (n=46) resulting into 376 significantly changing glycopeptides. (**Fig 5B**).

Several of the top changing downregulated glycoproteins (FC <0.7, p<0.05) contained complex (e.g. DKK3-N^204^ H5N4F1, CADH2-N^692^ H5N4A2F1) and hybrid glycans (e.g. DSC2-N^629^ H6N3A1F1) while among the top upregulated glycoproteins (FC>1.3, p<0.05) majority contained high mannose or paucimannose glycans (e.g. BNIP3-N^212^ H9N2, LAMP2-N^356^ H3N2F1, LAMP1-N^249^ H4N2F1, LAMC1-N^1439^ H7N2) or incomplete complex glycans (e.g., LAMP1-N^249^ H5N4F1, NPNT-N^273^ H5N4A1F1) (**Fig 5B, C)**.

Almost half of the most significantly changing glycopeptides (n=187) contained high mannose glycans, which is consistent with impaired ER glycosylation[61]^15^. In addition, several complex glycoproteins were significantly decreased, indicating GA glycosylation was similarly impaired (**Fig 5B**). These findings are in line with the previous observations that PGM1-deficiency results in abnormal glycosylation in both ER and GA[5,11], caused by the depletion of UDP-glucose and UDP-galactose[5] (**Fig 5A),** and confirm PGM1 deficiency also results in abnormal glycosylation in the heart. The most significantly changing glycopeptides belonged to ECM (eg. NPNT, CO5A2, EMIL1, ALCAM), and membrane/lysosomal trafficking (LAMP1, LAMP2) and only a few glycopeptides of metabolism/mitochondrial function (e.g., BNIP3, LRP1, SAP) (additional data). Oral galactose supplementation has been shown to effectively improve glycosylation in PGM1-CDG, however the heart involvement in PGM1-CDG is not treatable by galactose[3,5,12,14,62]. Considering the majority of the mitochondrial and sarcomeric proteins are not glycosylated, improvement in glycosylation by galactose would not have a direct effect on these proteins. These findings suggests that while glycosylation might be affected in the PGM1-deficient cardiomyocytes, disrupted sarcomere and mitochondrial dysfunction are the main drivers of galactose-resistant cardiomyopathy.

### Glucose flux is altered in PGM1-deficient iCM

To functionally confirm the deleterious effect of PGM1 deficiency on cardiac energy metabolism and mitochondrial function, we performed tracer metabolomics experiments. We incubated PGM1-deficient and CTR iCMs with 5.5mM (physiologic) ^13^C_6_-glucose, performed metabolite extraction and subjected the metabolites to targeted LC/MS analysis (**Fig 6A**). We then assessed metabolite abundances,^13^C_6_-glucose fractional contribution (FC, the percentage of metabolite labeled and isotopologues labeling (the number of carbons coming from ^13^C_6_-glucose in each metabolite). Previously, to assess the therapeutic effect of galactose in PGM1-deficient fibroblasts, we have performed tracer experiments in PGM1-deficient fibroblasts, using either ^13^C_6_ galactose or ^13^C_6_ glucose. Unfortunately, designing an experiment using ^13^C_6_ galactose in iCMs was not possible, as the medium used to differentiate iCMs naturally contains ^12^C_6_-galactose (approximately less than 0.5mM), which cannot be removed without changing the differentiation protocol. Nevertheless, even in the presence of 0.5 mM galactose in the medium, we found that PGM1-deficient iCMs clustered separately from the CTR as demonstrated by the heatmap (**Fig 6B**). The abundance of several metabolites was significantly altered compared to the CTR iCMs **(Fig 6C**). Unlike in PGM1-deficient fibroblasts, UDP-hexose (pool of UDP-galactose and UDP-glucose) in PGM1-deficient iCMs was not significantly decreased, likely due to the presence of galactose (**Fig 6B**). On the other hand, hexose (pool of glucose, galactose, mannose, fructose), erythrose-4-P, alanine, glutathione, propionyl-CoA and nucleotide-triphosphates (ATP, GTP, CTP, UTP) were among the most significantly decreased metabolites in PGM1-deficient iCMs (**Fig 6C, D**). To understand which pathways were the most affected in PGM1-deficient iCMs, we also performed pathway analysis using MetaboAnalyst[34]. The most significantly affected metabolic pathways included pyrimidine; glutathione; alanine, aspartate and glutamate metabolism as well as TCA cycle; arginine and pyruvate metabolism (**Fig 6 D, E**), all strongly linked to mitochondrial function. Finally, the fractional contribution (FC) and isotopologues labeling of ^13^C_6_-glucose showed profound metabolic rewiring in PGM1-deficient iCMs resulting in an altered glucose flux through metabolites of pentose phosphate pathway (PPP) nucleotide synthesis, and TCA cycle (**Fig 6 E, Sup Fig 2, 3**). These results show a metabolic rewiring in the PGM1-deficient iCMs resulting in impaired nucleotide synthesis and energy metabolism, corroborating our findings that PGM1 deficiency in cardiomyocytes leads to mitochondrial dysfunction and energy failure (**Fig 2, 3, 4**).

**Figure 6.**
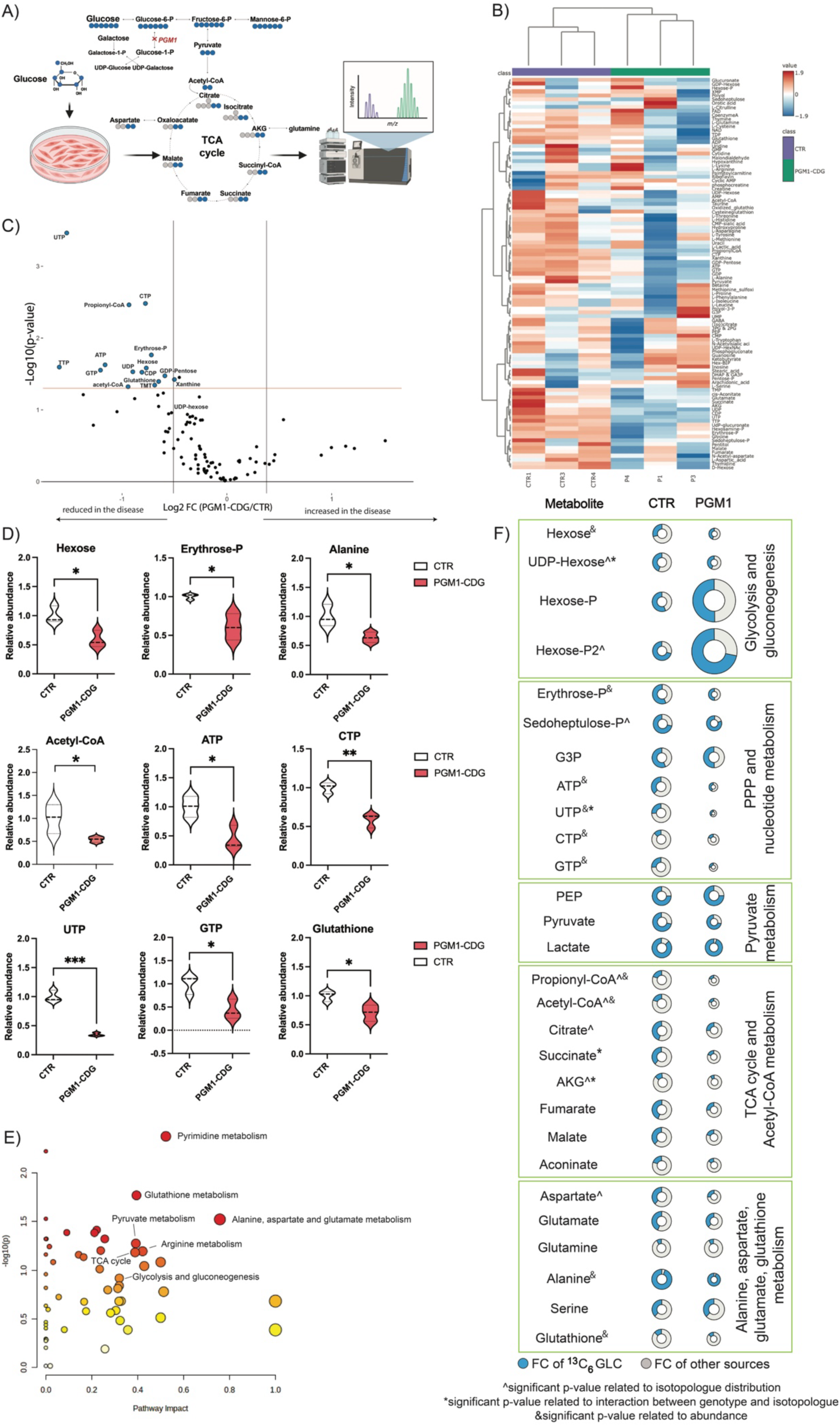
Glucose flux is altered in PGM1-deficient iCMs. **A)** Schematic representation of the tracer metabolomics methodology. **^13^C_6_-glucose is added to the medium not containing glucose (but containing ^12^C_6_-galactose). B)** Volcano plot showing the most significantly changing metabolites (FC< 0.7, p-value <0.05) in PGM1-deficient iCMs. **C)** Heatmap of all relatively quantified metabolites in CTR and PGM1-deficient iCMs. **D)** Violin plots showing the most significant metabolite changes in PGM1-deficient iCMs compared to CTR. Significance is indicated as * p<0.05, ** p<0.01, *** p<0.001. **E)** Pathway analysis with top identified changing pathways in PGM1-deficient iCMs. **F**) Schematic representation of the metabolic consequences of PGM1 deficiency and altered glucose flux in metabolites identified by tracer metabolomics analyses. Significant differences in isotopologues (positional) labeling of the metabolites were assessed by TWO-way ANOVA. Specific changes in positional labeling are represented in Sup Fig 2 and Sup Fig 3. Significant differences in labeling related to either genotype or interaction between genotype are indicated either **^** or **\*** respectively. The significant differences in the abundance of the metabolites are indicated with a **&**. PGM1 n=3, t=2-3; CTR n=3, t=2-3. Violin plots are shown with standard deviation (SD). Detailed statistical analysis can be found in the supplemental information.

### PGM1-deficient iCM have decreased basal and ATP-linked respiration

To further functionally validate the omics findings, and to specifically assess the respiration in PGM1-CDG cardiomyocytes we performed respiration studies using Seahorse XFe96 (Agilent) (see methods). PGM1-deficient iCMs displayed reduced oxygen consumption rate (OCR) relative to cell number (proxy for DNA and protein content) **Fig 7A**) at all time points of the mito stress test assay, while no difference in proton efflux rate (PER) was observed (**Fig 7D**). PGM1-CDG iCMs displayed significantly reduced basal respiration, ATP-associated respiration, proton leak, maximal respiration, and spare OCR relative to controls (**Fig 7A, B**). The metabolic map of resting OCR and PER showed PGM1 iCMs clustered separately from the controls, characterized by greatly reduced OCR and only moderately reduced PER (**Fig 7C**). While basal PER was not significantly reduced in PGM1-CDG iCMs, spare glycolytic capacity (the change in PER following oligomycin administration) as a percentage of basal PER was highly significantly reduced relative to controls(**Fig 7E**). Based on the OCR and PER data derived using the Cell Mito Stress Test kit, pmol of ATP produced through oxidative phosphorylation (OxPhos) and glycolysis were calculated according to the manufacturer instructions. PGM1-CDG iCMs displayed a highly significantly reduced total ATP production, as well as a highly significantly reduced OxPhos-derived ATP production (**Fig 7F**).

**Figure 7.**
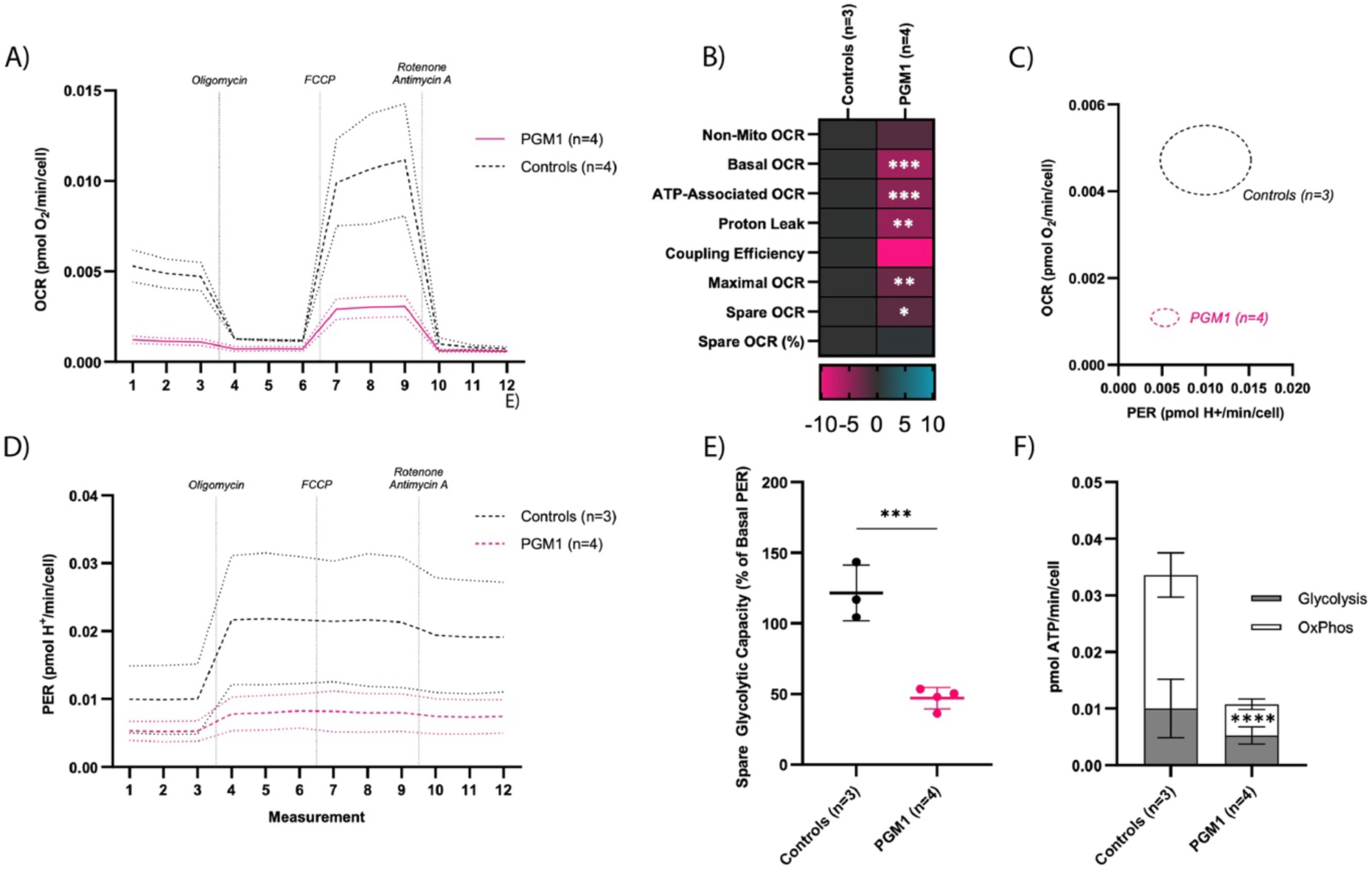
PGM1-deficient iCM show decreased basal, maximal and ATP-linked respiration. Mitochondrial stress test normalized to cell count. A) Oxymetry plot displaying Oxygen Consumption Rate (OCR) of PGM1 and Control iCMs (mean +/- SD) during Mito Stress Test Assay. B) Heat map of Mito Stress Test OCR readouts, representing SDs from the mean of the controls. C) Metabolic map displaying Proton Efflux Rate (PER) on the X axis and OCR on the Y axis. Circles represent SD of the means of each group. D) Proton efflux rate (PER) plot displaying PER of PGM1 and Control iCMs (mean +/- SD) during Mito Stress Test Assay. E) Spare glycolytic capacity. F) Bar plot displaying sources of ATP. B) Mitochondrial stress test normalized to CS. The P values are indicated as significant * (<0.05), ** (<0.01), or *** (<0.001). (PGM1 n=4, t=3 CTR n=3, t=3)

As the differences in respiration can also be driven by the changes in mitochondrial mass, we also normalized the data to citrate synthase (CS) activity, which is a proxy readout for mitochondrial mass. Normalized to CS, the basal, maximal and ATP-linked respiration were significantly decreased in PGM1-deficient iCMs **(Additional Fig 4A-F**), as was the spare glycolytic capacity. There was no significant difference in CS activity between PGM1-CDG and CTR iCMs (**Additional Fig 4G**). These findings indicate that PGM1-deficient iCMs exhibit severely reduced mitochondrial respiration, primarily driven by alterations in mitochondrial ATP-linked respiration.

### *In silico* drug repurposing reveals potential drug candidates targeting cardiac development and mitochondrial function

Though tremendous progress has been made in treating mitochondrial dysfunction in mitochondrial disease[63], unfortunately there are still no approved FDA treatments for mitochondrial disease. Therefore, treating mitochondrial dysfunction remains a challenge. To identify potential novel cardiac-specific treatment strategies in PGM1-CDG, we used Ensemble of Multiple Drug Repositioning Approaches (EMUDRA)[39] on our untargeted proteomics data. EMUDRA compares the effects of ∼1300 FDA approved small molecules and predicts whether these molecules can effectively normalize log2foldchange values (see methods for details). EMUDRA identified several drugs that are predicted to downregulate processes that are upregulated in PGM1-deficient iCMs, and were also reflected in the GSEA results, such as vesicle organization, ER to Golgi transport, protein targeting to mitochondria (**Fig 8A**). In addition, EMUDRA identified several drugs that are predicted to upregulate processes that are downregulated in PGM1-deficient iCMs, such as mitochondrial genome maintenance, pyruvate metabolism, cardiac muscle tissue development, muscle structure and cell development, and cell communication involved in cardiac conduction (**Fig 8B**). Several drugs were shown to target both upregulated and downregulated processes. These included for example NSAIDs such as zomepirac, chlorzoxazone, a muscle relaxant, betaxolol, β1selective (cardioselective) adrenergic antagonist, and compound 5109870, which was previously shown to increase PGM1 expression *in vitro* (**Fig 8A, B**).

**Figure 8.**
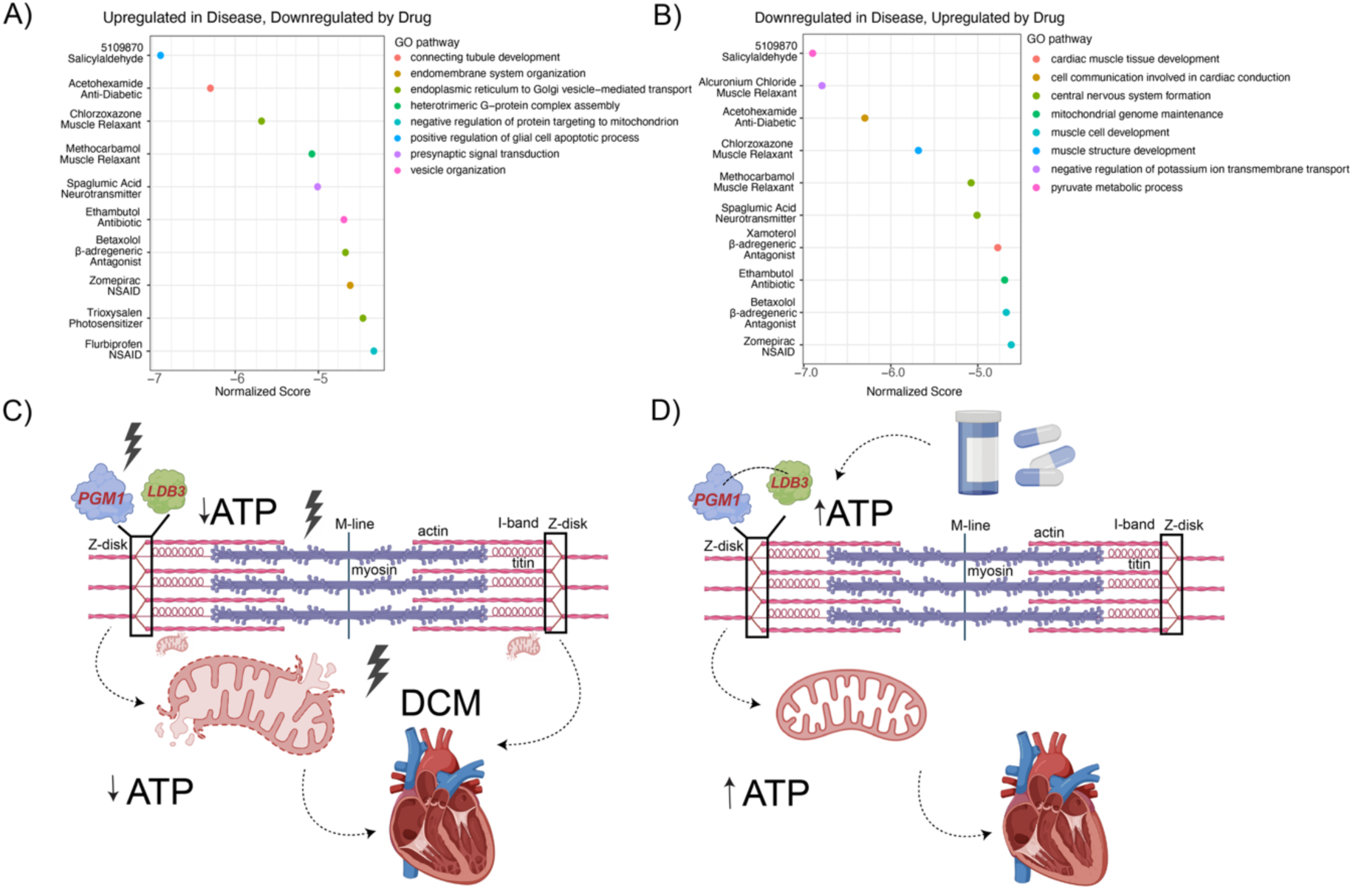
Exploring therapeutic options for cardiac presentation in PGM1 deficiency. **A)** EMUDRA identified drugs that are predicted to upregulate the processes that are downregulated in PGM1-deficient iCMs based on Differentially Expressed Gene analysis (log2foldchange <0.25, p-value <0.05). B) EMUDRA identified drugs that are predicted to downregulate the processes that are upregulated in PGM1-deficient iCMs based on Differentially Expressed Gene analysis (absolute log2foldchange >0.25, p-value <0.05). The top-11 drug candidates are shown for both analysis. The Drug Class is written below the CMAP drug. Normalized enrichment Scores indicate the potential of the drug to normalize the phenotype, and lower scores are associated with higher potential of normalizing aberrant log2foldchange values. C) potential pathomechanism linking PGM1 deficiency to sarcomere disruption and mitochondrial dysfunction, eventually resulting in cardiac failure in PGM1-CDG, D) potential therapeutic mechanism in PGM1 deficiency targeting sarcomeric proteins resulting in improved cardiac function.

In conclusion, our findings suggest that the PGM1 plays a crucial role in cardiac architecture. The PGM1 deficiency possibly results in the destabilization of Z-disk through impaired LDB3 binding, and subsequent mitochondrial dysfunction (**Fig 8C**). Targeting PGM1 deficiency by one of the drugs identified by EMUDRA could therefore result in the restoration of cardiac machinery and mitochondrial function (**Fig 8D**), and a better clinical outcome for patients with PGM1-deficiency.

## Discussion

PGM1 deficiency is associated with severe, often lethal cardiac complications. While its role in glycogen metabolism and glycosylation is well established, the specific function of PGM1 in the heart remains unclear. To address this gap, we generated and characterized PGM1-deficient iCMs derived from individuals with pathogenic *PGM1* variants (**Fig 1B-E**). These iCMs exhibited reduced beating capacity and contractility (**Fig 1F-K**). To uncover the cardiac-specific pathomechanisms underlying PGM1 deficiency, we conducted comprehensive multi-omics analyses to identify molecular drivers of the observed cardiomyocyte dysfunction.

Proteomic analysis revealed a significant decrease in PGM1 expression, accompanied by upregulation of extracellular matrix (ECM) proteins and marked downregulation of key proteins involved in sarcomere integrity, cardiac function, and mitochondrial activity (**Fig 2**). Given the depletion of cardiac and sarcomeric proteins, we hypothesized that PGM1 may have a structural role in maintaining cardiac architecture.^17^ PGM1 has previously been shown to associate with the Z-disk protein Zasp/Cypher (LDB3)[17] Pathogenic variants in *LDB3* have been shown to disrupt PGM1-LDB3 binding under stress conditions, leading to defective Z-disk formation and the development of dilated cardiomyopathy[17]. As *LDB3* contains cardiac-specific exons that may serve as potential binding sites for PGM1, we hypothesized that *PGM1* mutations impair PGM1-LDB3 binding, leading to Z-disk destabilization and exacerbating the biochemical consequences of PGM1 deficiency in the heart. Supporting this hypothesis, we observed reduced LDB3 levels in PGM1-deficient iCMs, along with decreased expression of other key Z-disk components such as ACTN2 and MYOZ2 (**Fig 2**, Additional Data). *In silico* binding studies using AlphaFold3[37] further implicated exon 4 of *LDB3* as a potential interaction site for PGM1 (**Fig. 2C-F**). The *in silico* predictions were then confirmed by *in vitro* binding experiments, specifically showing PGM1 binds to exon 4 of LDB3 (**Fig 2G**). These findings suggest that PGM1 (or specifically its cardiac isoform PGM1-2) bind to LDB3 via exon 4 and that PGM1 deficiency disrupts Z-disk integrity, contributing to cardiac dysfunction. Since exon 4 is cardiac-specific, this mechanism could also explain why the heart fails to respond to galactose treatment, unlike other organs where alternative *LDB3* isoforms are present and sarcomeric structure is less critical.

The sarcomere Z-disk and mitochondrial function in the heart are closely interconnected[48,57,58,64,65]. Intermyofibrillar scaffolds within the Z-disk serve as anchors for thin filaments, transmitting force along myofibrils while regulating their length changes and preventing membrane damage[57,65,66]. These scaffolds are also essential for mitochondrial localization, as mitochondria are positioned at the I-b and. Disruptions in myofibrillar structure often lead to abnormal mitochondrial distribution, and myofibrillar myopathies, including *LDB3* deficiency, are frequently associated with mitochondrial dysfunction[57,65,66]. Similarly, Z-disk disarray and swollen mitochondria were previously shown in PGM1 cardiac specific KO mouse model showed and in the heart of the patient with PGM1-CDG[19]. To assess the extent of mitochondrial abnormalities in PGM1-deficient iCMs, we performed MitoCart analysis. MitoCarta analysis revealed a significant depletion of mitochondrial proteins, including components of complexes I–V of the electron transport chain, as well as proteins involved in mitochondrial biogenesis and quality control (**Fig 3**), corroborating the previous findings and suggesting a mechanistic link between PGM1 deficiency, Z-disk instability, and mitochondrial dysfunction.

Further investigation using Ingenuity Pathway Analysis (IPA) highlighted several affected pathways, including mitochondrial function, ECM regulation, protein sorting, and ER-to-Golgi transport pathways (**Fig 4**). Toxicity analysis revealed that the PGM1-deficient proteome overlaps with signatures of cardiac arrhythmia, dilation, and enlargement—common cardiac manifestations in PGM1-CDG (**Fig 4B**). Both mitochondrial and heart disorders were strongly associated with the PGM1-deficient proteome (**Fig 4C**), and the GSEA confirmed mitochondrial dysfunction, cell cycle dysregulation, and ECM remodeling as affected pathways in PGM1-deficient iCMs (**Fig.4D-F**).

All significantly altered processes identified by pathway analysis in PGM1 iCMs, ECM remodeling, and mitochondrial dysfunction—are closely linked to cardiac failure including dilated cardiomyopathy[58]. Both ECM upregulation[59,67] and mitochondrial and OXPHOS dysfunctions have been linked to dilated cardiomyopathy[58,68], the most common cardiac manifestation in PGM1-CDG.^44^ Previous reports of abnormal mitochondrial architecture[19] and fibrosis[9] in individuals with PGM1-CDG[19] further support the connection between these processes and cardiac pathology.

To dissect the contribution of glycosylation abnormalities in the PGM1-cardiomyophaty, we conducted glycoproteomics analysis in our iCMs. PGM1-deficient iCMs exhibited significant alterations in glycoprotein profiles (**Fig. 5**), with upregulated glycoproteins predominantly containing high mannose/paucimannose glycans, while those with complex glycans were markedly downregulated (**Fig. 5A-C**). This pattern closely resembled the glycosylation defects observed in the blood of PGM1-CDG patients, where high mannose and truncated glycans are elevated, while complex glycans are reduced[11,13]. Notably, the most affected glycoproteins were associated with extracellular matrix remodeling, membrane trafficking, and only a few pertaining to metabolism and mitochondrial function (**Fig. 5C**). Considering glycosylation abnormalities are treatable by galactose[11,14], we hypothesize that ECM related abnormalities and membrane trafficking proteins, which are heavily glycosylated, would respond to galactose. On the other hand, the majority of sarcomeric and mitochondrial proteins are not glycosylated, and their expression independent of glycosylation, further highlighting sarcomere (and mitochondrial) disruption as drivers of therapy-resistant PGM1-deficient cardiomyopathy.

Energy failure in PGM1-deficient iCMs was further supported by our ^13^C_6_-glucose tracer studies. While UDP-hexose levels were not significantly reduced, likely due to the presence of galactose in the culture medium, the most disrupted pathways were nucleotide metabolism and the TCA cycle (**Fig. 6**). Tracer analysis revealed altered glucose flux through the pentose phosphate pathway (PPP), nucleotide synthesis, and the TCA cycle, highlighting metabolic shifts in PGM1-deficient iCMs (**Fig. 6E, Sup. Fig. 2,3**). As the TCA cycle is directly linked to oxidative phosphorylation (OXPHOS) and mitochondrial function, disruptions in nucleotide synthesis have broader implications beyond energy metabolism, affecting cell cycle regulation and cardiac function. Notably, mitochondrial dysfunction is known to impair de novo pyrimidine synthesis through dihydroorotate dehydrogenase (DHODH)[69], a mitochondria-coupled enzyme. Additionally, previous studies have shown that electron transport chain (ETC) dysfunction suppresses *de novo* purine synthesis while upregulating the purine salvage pathway[70]. In the heart, nucleotides play a critical role, as ATP drives essential cellular and mechanical processes, including the actin-myosin bridging cycle. Depletion of nucleotides and ATP has been implicated in impaired cardiac mechanics and cardiac failure[71,72]. Consistent with this, mitochondrial dysfunction and ATP depletion in PGM1-deficient iCMs were further validated through Seahorse respiration studies (**Fig 7, Sup. Fig. 4**).

Identifying therapeutic strategies with immediate clinical translation is crucial for PGM1-CDG patients with cardiac complications. To address this, we performed in silico drug repurposing using the Ensemble of Multiple Drug Repositioning Approaches (EMUDRA)[39] to identify potential treatments for galactose-resistant cardiac dysfunction in PGM1-CDG (**Fig 8**). EMUDRA analysis identified several promising drug candidates, including antibiotics, HDAC inhibitors, and NSAIDs, which target pathways disrupted in PGM1-deficient iCMs. Among them, zomepirac, an NSAID, was predicted to downregulate ER-to-Golgi vesicle transport pathways while enhancing processes related to cardiac and muscle cell development as well as mitochondrial maintenance (**Fig 8A**). However, due to the potential risks of long-term NSAID use, alternative candidates may provide safer therapeutic options. Further *in vitro* testing in PGM1-deficient iCMs could help evaluate the effects of these candidates on mitochondrial function, though this falls beyond the scope of the present study.

In conclusion, our study provides new insights into the cardiac-specific mechanisms of PGM1 deficiency, revealing its impact on sarcomere integrity, mitochondrial function, energy metabolism, and ECM regulation in cardiomyocytes. Our data indicate that PGM1 deficiency is associated with disrupted LDB3 interactions, altered sarcomeric structure, and impaired mitochondrial function, contributing to therapy-resistant cardiac dysfunction (**Fig 8C**). This is consistent with clinical observations that galactose treatment, which improves systemic symptoms, does not ameliorate cardiac or skeletal muscle phenotypes in individuals with PGM1-CDG [3,5,11,12]. Finally, these findings support the rationale for investigating mitochondrial-targeted therapies as a potential approach to address cardiac involvement in PGM1-CDG (**Fig 8D**).

## Conclusions

Here, we report that PGM1 plays a critical role in maintaining cardiac structure and function. Our findings suggest that PGM1 deficiency may destabilize the Z-disk by disrupting its interaction with LDB3, ultimately leading to mitochondrial dysfunction (**Fig 8C**). Furthermore, we show that PGM1-deficient iCMs exhibit glycosylation abnormalities, primarily affecting glycoproteins involved in extracellular matrix organization and membrane trafficking, which might be targeted by galactose. On the other hand, sarcomere and mitochondrial proteins are not glycosylated, and their status not dependent on glycosylation. These results highlight mitochondrial dysfunction as a potential therapeutic target to improve cardiac performance and clinical outcomes in individuals with PGM1-CDG (**Fig 8D**)

### Limitations of the study

The main aim of this study was to assess the role of PGM1 in cardiomyocytes, with a special focus on Z-disk and mitochondria. The pathomechanism leading to glycosylation abnormalities in PGM1 deficiency as well as the ability of galactose to correct glycosylation abnormalities in PGM1-CDG has been extensively studied[5,11,14]. While we assessed the glycosylation abnormalities in PGM1-deficienct iCMs, we have not explicitly performed galactose treatment experiments in iCMs, and we can only speculate about the effect of galactose on these abnormalities. In addition, the medium used to culture iCMs contains galactose (B27 supplement). While the galactose concentration in the iCM medium is lower (0.5mM) than levels known to significantly improve glycosylation in fibroblasts (2mM)[5], the presence of galactose could have mitigated the glycosylation abnormalities in PGM1 iCMs *in vitro*. Nevertheless, regardless of the presence of galactose in the medium, PGM1 iCMs showed extensive sarcomere and mitochondrial-related abnormalities further suggesting these proteins are not responding to galactose.

## Declarations

### Ethics

Informed research consent was obtained from all patients included in this study. Deidentified, residual samples from PGM1-CDG affected individual were used to create hiPSCs. hiPSC cell lines were derived from fibroblasts previously collected as part of clinical management and evaluation and stored in the Frontiers of Congenital Disorders of Glycosylation (FCDGC) biobank at Mayo Clinic (Mayo Clinic *IRB: 16-004682*). Additional healthy fibroblasts were obtained from Coriell institute (GM01651, GM08399). Two hiPSC cell lines (1363, 8858) were a gift from Dr. Sergiu Pasca and previously reported[20]. Demographic and genetic information of the PGM1-CDG individuals whose cell lines were used is given in **Table 1**.

### Materials availability

All reagents are commercially available. Apart from the hiPSC lines, this study did not generate unique new reagents. Further information and requests for resources and/or reagents can be directed to the lead contact.

### Data availability

- Deidentified metabolomics data have been deposited at National Metabolomics Data Repository (NMDR) Metabolomics workbench and are publicly available as of the date of publication.
- Deidentified mass spectrometry glycoproteomics and proteomics data have been deposited to the ProteomeXchange Consortium via the PRIDE partner repository.
- This study does not report a novel code.
- All the data reported in this paper such as values and complete statistical analysis used to create graphs in this paper are available as the additional data.
- All data and algorithms generated in this study can be requested from the corresponding author (Tamas Kozicz). Any additional information needed to analyze the data can be received from the lead contact upon request.

### Conflict of interest

Authors have no conflicts of interest to declare.

### Author contributions

Study conceptualization: EM, KL, TK; methodology: ANL, AP, EM, GP, LJB, NPS, RH, RB, SR, TK; investigation: ANL, GP, IB, IM, IS, LJB, NPS, RH RB, RS, SR, SV; formal analysis: EM, GP, RB, SR, TK; resources: AP, EM, KL, TK; data curation: AP, GP, IM, RB, SR, TK; writing original draft: EM, SR, TK; writing-review and editing: ANL, AP, BB, EM, GP, IB, IM, IS, LJB, NPS, RH, RB, RS, SR, KL, TK; visualization: GP, IM, LJB, SR; supervision: AP, EM, TK; project administration: AP, EM, KL, TK; funding acquisition: AP, EM, KL, TK. All authors have read and approved the manuscript.

### Funding

This work was supported by 1U54NS115198*-01* from the National Institute of Neurological Diseases and Stroke (NINDS), the National Center for Advancing Translational Sciences (NCATS), the National Institute of Child Health and Human Development (NICHD) and the Rare Disorders Consortium Disease Network (RDCRN).

## Acknowledgements

The authors would like to thank Samantha Hamrick, MSc. and Alexander Kreymerman, PhD. for her help with the iCardiomyocyte protocol set up.

## Notes

### Competing Interest Statement

The authors have declared no competing interest.

