## Supplementary information and figures for "PGM1 deficiency disrupts sarcomere and mitochondrial function in a stem-cell cardiomyocyte model"

### Supplementary (additional) material

Additional material contains figures related to the work in the main text. Supplemenary workbook contains raw data generated from each omics analysis and statistical analysis.

### Additional Figures

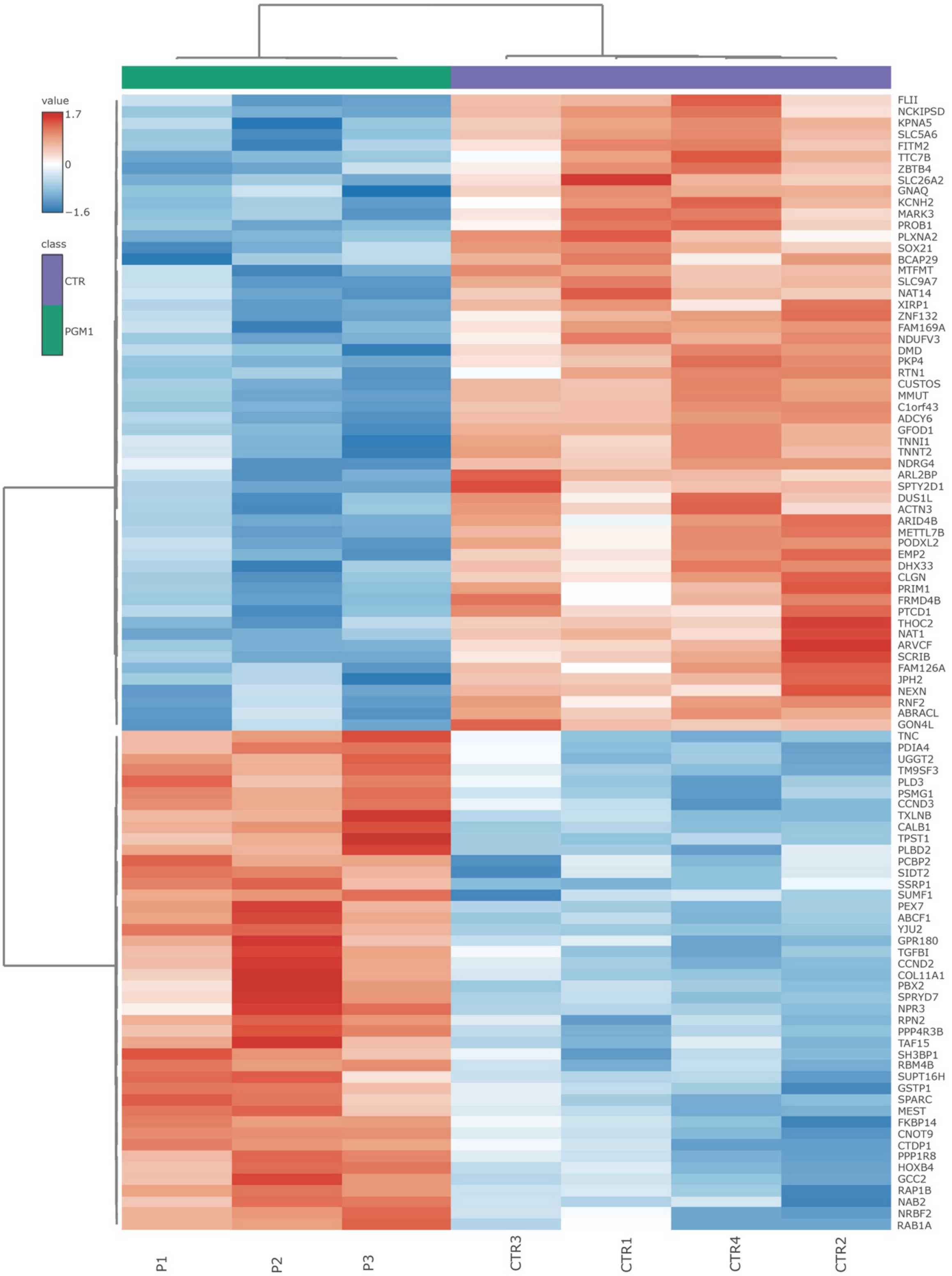


**Additional Figure 1.** Heatmap of top 100 significantly (p-value < 0.05) upregulated and downregulated proteins in PGM1-deficient iCM PGM1 n=3, t=1; CTR n=4, t=1.

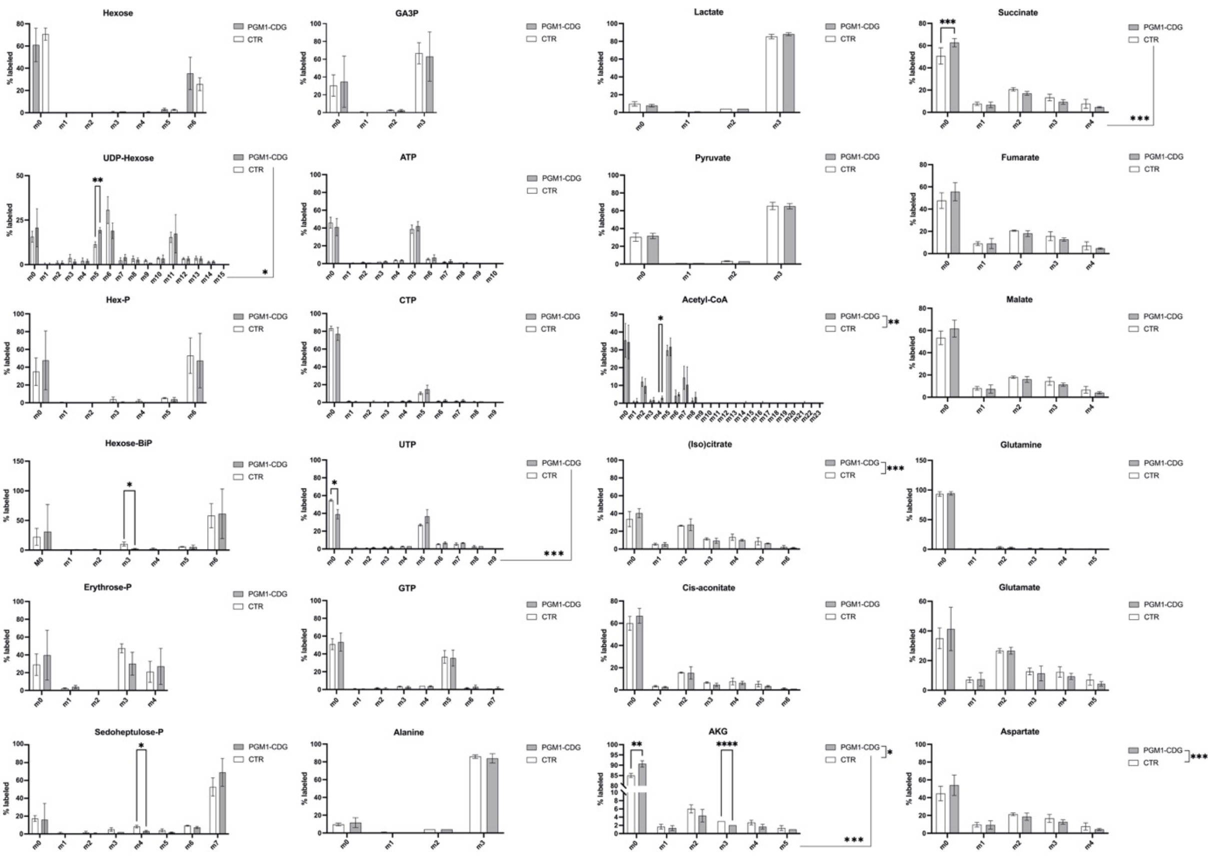


**Additional Figure 2.** Isotopologue distribution (positional labeling) of $^{13}C_6$-glucose in specific metabolites belonging to glucose and galactose metabolism, pentose phosphate pathway, nucleotide-phosphates, TCA cycle and glutamine metabolism. m(0-n) represents the number of carbons labeled by $^{13}C_6$-glucose, where n=number of carbons present in the metabolite. Two-way ANOVA and multiple comparisons with Šidak correction was performed. Significant p-value is indicated in * (* p<0.05; ** p< 0.01; *** p<0.001). (PGM1 n=3; t=2-3; CTR n=3, t=2-3). Detailed statistical analysis is provided in Additional data.

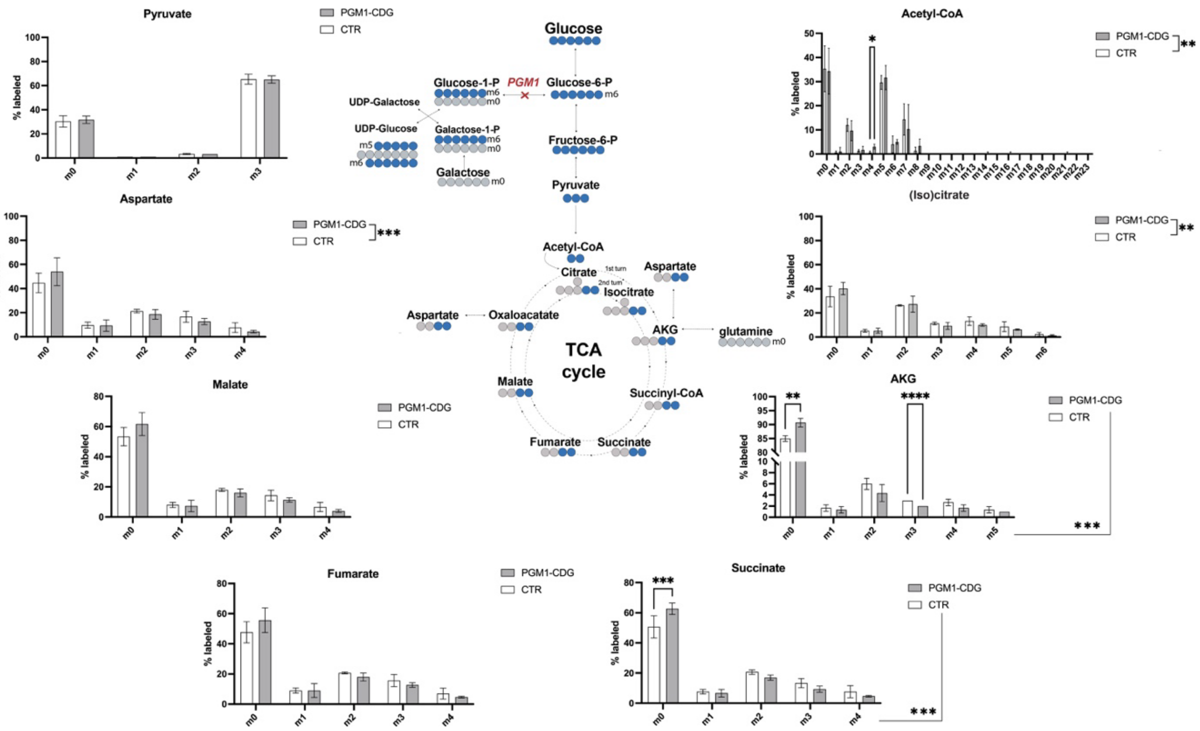


**Additional Figure 3.** Isotopologue distribution (positional labeling) of $^{13}C_6$-glucose in related to the TCA cycle. m(0-n) represents the number of carbons labeled by $^{13}C_6$-glucose, where n=number of carbons present in the metabolite. Two-way ANOVA and multiple comparisons with Šidak correction was performed. Significant p-value is indicated in * (* p<0.05; ** p< 0.01; *** p<0.001). (PGM1 n=3; t=2-3; CTR n=3, t=2-3). Detailed statistical analysis is provided in Additional data.

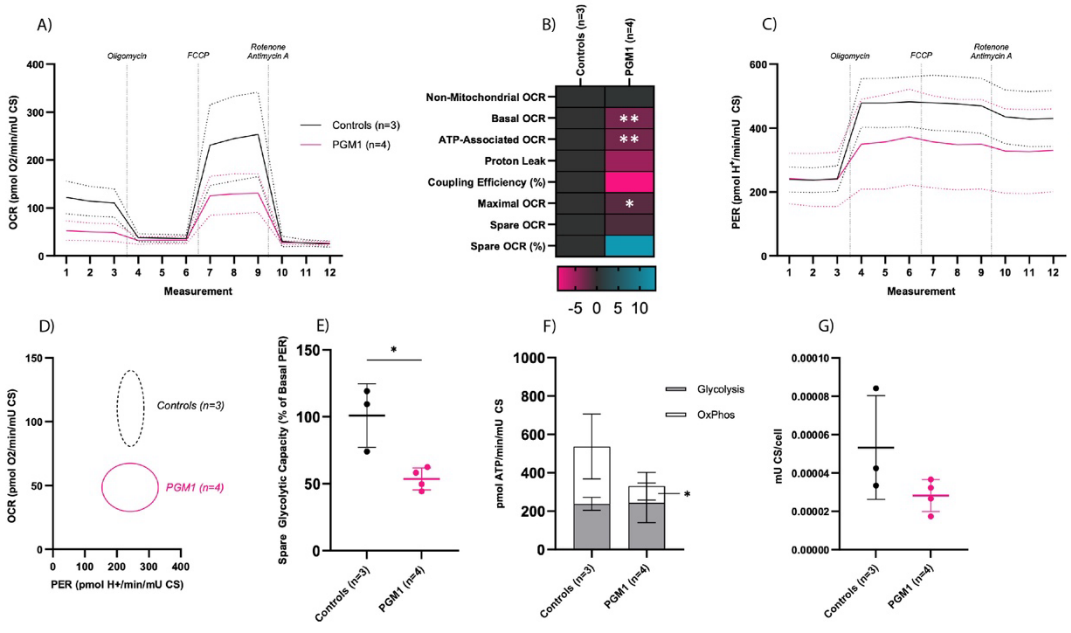


**Additional Figure 4.** Mitochondrial stress test normalized to Citrate Synthase (CS). A) Oxymetry plot displaying Oxygen Consumption Rate (OCR) of PGM1 and Control iCMs (mean +/- SD) during Mito Stress Test Assay. B) Heat map of Mito Stress Test OCR readouts, representing SDs from the mean of the controls. C) Proton efflux rate (PER) plot displaying PER of PGM1 and Control iCMs (mean +/- SD) during Mito Stress Test Assay. D) Metabolic map displaying Proton Efflux Rate (PER) on the X axis and OCR on the Y axis. Circles represent SD of the means of each group. E) Spare glycolytic capacity. F) Bar plot displaying sources of ATP. G) Citrate Synthase (CS) activity normalized to cell count. The P values are indicated as significant * (<0.05), ** (<0.01), or *** (<0.001). (PGM1 n=4, t=3 CTR n=3, t=3).
